# Illuminating microbial mat assembly: Cyanobacteria and Chloroflexota cooperate to structure light-responsive biofilms

**DOI:** 10.1101/2024.07.24.605005

**Authors:** Freddy Bunbury, Carlos Rivas, Victoria Calatrava, Andrey Malkovskiy, Lydia-Marie Joubert, Amar D. Parvate, James E. Evans, Arthur Grossman, Devaki Bhaya

**Affiliations:** Department of Ecology and Evolution, The University of Chicago, Chicago, IL 60637, USA; Department of Biosphere Sciences and Engineering, Carnegie Institution for Science, 260 Panama St., Stanford, CA 94305, USA; SLAC National Accelerator Laboratory, Division of CryoEM and Bioimaging, Menlo Park, CA 94025, USA; Pacific Northwest National Laboratory, Environmental Molecular Sciences Laboratory, Richland, WA, USA; Washington State University Pullman, School of Biological Sciences, Pullman, WA, USA; Stanford University, Biology Department, Stanford, CA 94305, USA

## Abstract

Microbial mats are stratified communities often dominated by unicellular and filamentous phototrophs within an exopolymer matrix. It is challenging to quantify the dynamic responses of community members *in situ* as they experience steep gradients and rapid fluctuations of light. To address this, we developed a binary consortium using two representative isolates from hot spring mats: the unicellular oxygenic phototrophic cyanobacterium *Synechococcus* OS-B’ (Syn OS-B’) and the filamentous anoxygenic phototroph *Chloroflexus* MS-CIW-1 (Chfl MS-1). We quantified the motility of individual cells and entire colonies and demonstrated that Chfl MS-1 formed bundles of filaments that moved in all directions with no directional bias to light. Syn OS- B’ was slightly less motile but exhibited positive phototaxis. This binary consortium displayed cooperative behavior by moving further than either species alone and formed ordered arrays where both species aligned with the light source. No cooperative motility was observed when a non-motile *pilB* mutant of Syn OS-B’ was used instead of Syn OS-B’. The binary consortium also produced more adherent biofilm than individual species, consistent with the close interspecies association revealed by electron microscopy. We propose that cyanobacteria and Chloroflexota cooperate in forming natural microbial mats, by colonizing new niches and building robust biofilms.

**Significance:** Microbial mats are dense, layered communities with ancient origins and widespread occurrence, but how they assemble is not well understood. To investigate how microbial motility, physical interactions, and responses to light affect mat assembly, we developed a binary consortium from representative hot spring mat isolates. Individually, the Cyanobacteria and Chloroflexota isolates displayed significant differences in motility and biofilm formation. When combined, the consortium exhibited enhanced motility towards light and formed more robust biofilms. This model consortium approach complements *in situ* studies by directly testing the role of motility and physical cooperation in shaping microbial mats, and could inform biofilm applications in industrial settings.

## Introduction

Stratified microbial mats dominated by phototrophs are examples of complex biofilms that often thrive in extreme environments such as hot springs, desert crusts, and hypersaline lagoons [1]. These dense biofilms experience light fluctuations due to changing cloud cover and more predictable diel oscillations: increasing sunlight in the morning drives oxygenic photosynthesis, which increases O2 concentrations, decreases CO2, and increases pH; the opposite occurs in the afternoon [2], [3]. Microbes can respond to changing conditions by acclimating or moving to more propitious environments by tracking chemical gradients (chemotaxis and aerotaxis) or light direction (phototaxis) [4], [5]. Studying biofilm development in the lab has led to a distinction in bacterial life stages between a motile swarming phase vs a biofilm phase where cells are encased in a matrix and non-motile [6]. However, other studies have demonstrated that surface-based motility can be important in determining biofilm structure [7], [8], that whole biofilm-like microcolonies can be motile [9], and that the same genes can control both biofilm formation and motility [10]. These observations suggest that bacteria may be quite motile within biofilms and microbial mats, which has consequences for how nutrients [11] and DNA [12] are transferred between species, and how the community responds to phage attack [13] or abiotic stress [14].

Several species of Cyanobacteriota (hereafter cyanobacteria) and Chloroflexota are abundant in hot spring microbial mats [15], core contributors to community metabolite fluxes [16], and thought to be the main determinants of mat architecture [17]. Vertical distributions of cyanobacteria in mats vary over the diel cycle, suggesting migration in response to changing light, pH, or oxygen conditions [18], [19]. Laboratory experiments have shown that filamentous and unicellular cyanobacteria can use Type IV pili (T4P) to move directionally in response to light gradients by phototaxis [20], [21]. Chloroflexota isolates move by gliding motility [22] possibly driven by tight-adherence pili [23]. How these species contribute or interact with each other to form biofilms is relatively unknown.

The current understanding of microbial mat structure is based mostly on examining in situ snapshots of species depth profiles [19], [24], [25], [26], while a few laboratory experiments have examined the motility of individual species [22], [27]. We used the cyanobacterium *Synechococcus* sp. JA-2-3B′a(2-13) [28], originally isolated from Octopus spring in Yellowstone National Park (YNP) (recently proposed to be renamed *Thermostichus sp. JA-2-3Ba*) [29], and a member of the Chloroflexota, Chfl MS-1 [30], from the nearby Mushroom spring. Both species are abundant in several hot springs in YNP [31] and their motility, or that of close relatives, has been individually characterized to some extent [22], [27], [32]. We reasoned that developing a binary consortium composed of these unicellular and filamentous species could allow us to examine the role of light and physical interactions in structuring “mat-like” biofilms.

We found that the binary consortium of Syn OS-B’ and Chfl MS-1 exhibited greater movement towards the light than either species individually. Replacing the wild-type Syn OS-B’ cells with a non-motile *ΔpilB* mutant, completely abrogated the cooperative effect suggesting that Syn OS- B’ motility was essential. Microscopy of the binary consortium revealed a physical association and parallel arrangement of Syn OS-B’ and Chfl MS-1 cells. Chfl MS-1 formed a more adherent biofilm than Syn OS-B’, and biofilm formation was increased further when the species were mixed. Inspection of binary consortia by electron microscopy revealed intercellular attachments, potentially facilitated by the secretion of extracellular polymeric substances (EPS). These binary consortia recapitulate some features of natural microbial mats and could be further developed to understand how mats are structured by dynamic light environments.

## Results

### Syn OS-B’ and Chfl MS-1 are both motile on surfaces, but only Syn OS-B’ is phototactic

Syn OS-B’ grows as slightly curved rods 5-10 µm long and 2 µm wide (Figure S1A), while Chfl MS-1 grows as an unbranched filament >100 µm long and 600 nm wide (Figure S1B).

Transmission electron microscopy (TEM) of uranyl-acetate-stained whole cells revealed T4P located near the cell poles in Syn OS-B’ (Figure S1C), and presumptive tight-adherence pili located near the septa between cells in a Chfl MS-1 filament (Figure S1D). Syn OS-B’ and Chfl MS-1 contained phycocyanin and bacteriochlorophyll c pigments, respectively (Figure S2), allowing them to be distinguished at a macroscopic scale.

To investigate Syn OS-B’ and Chfl MS-1 motility, both species were grown separately to late log-phase in DH10M and PE medium, respectively, plated on DH10 solidified with 0.4% agarose at an adjusted OD600=2, and incubated under either directional white light (60 µmol photons/m^2^/s) (Figure S3) or in darkness for 4 days. Fluorescence scanning measurements showed Syn OS-B’ (Figure 1A) and Chfl MS-1 (Figure 1G) have different collective motility patterns. Syn OS-B’ moves towards the light while Chfl MS-1 moves in all directions with no apparent bias. Syn OS-B’ increased its colony area only under directional light (Figure 1B), while Chfl MS-1 increased its colony area in both the light and dark (Figure 1H). Syn OS-B’ cells were phototactic, forming parallel finger-like projections towards the light so that the center of the colony shifted towards the light by 2 mm (Figure 1C, Video S1), as described in greater detail in [32]. Chfl MS-1 also produced collective projections composed of filament bundles, which formed more rapidly than those of Syn OS-B’ but with no directional bias (Figure 1I, Video S2).

**Figure 1.**
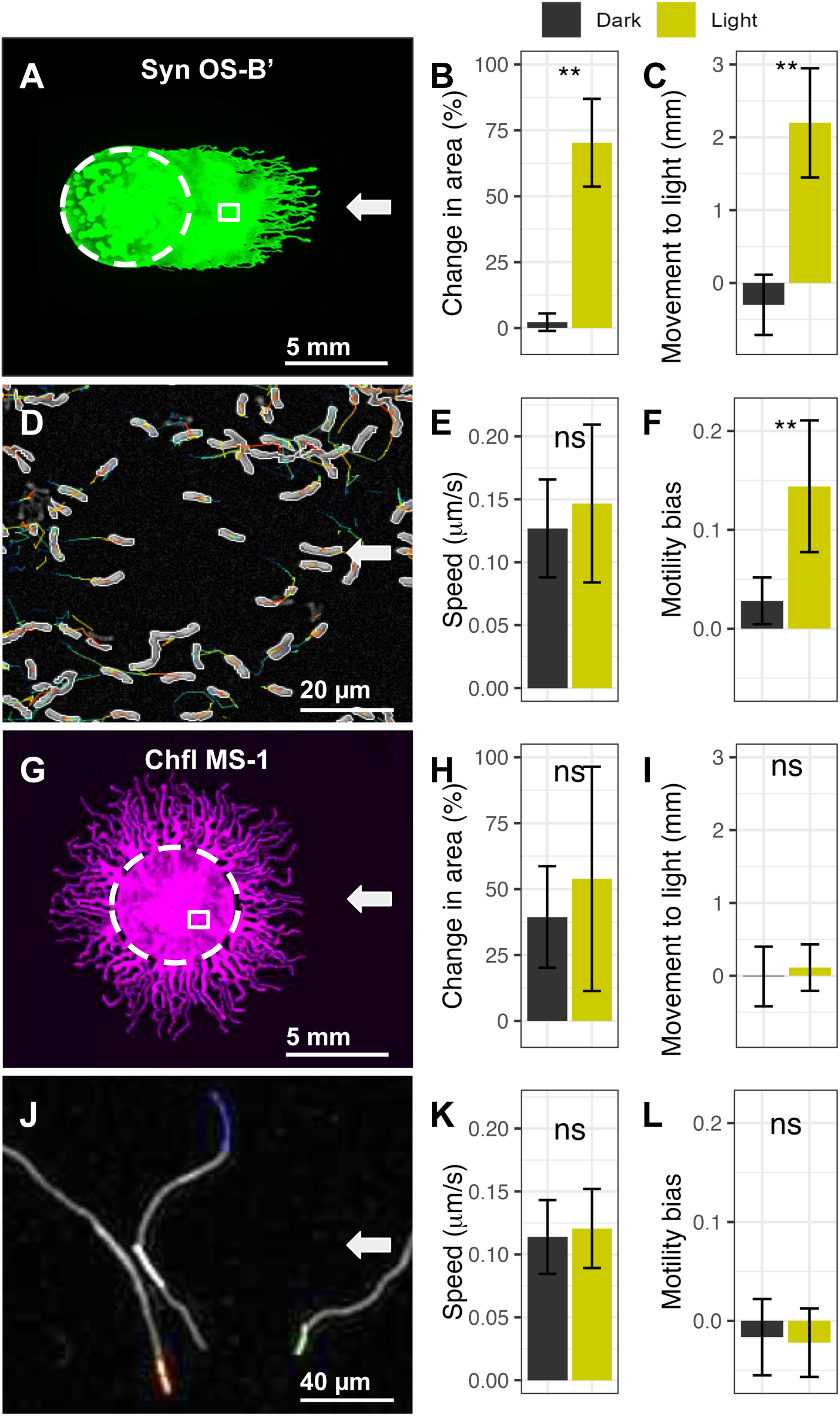
Syn OS-B’ and Chfl MS-1 are both motile, but only Syn OS-B’ is phototactic. **(A,G)** Fluorescence scan (ex. 658 nm / em. 710 nm) of a Syn OS-B’ colony after 4 days of movement under directional white light, and fluorescence scan (ex. 784 nm / em. 832 nm) of a Chfl MS-1 colony after 4 days of the same conditions. White arrows indicate light direction, dashed circles indicate initial colony/droplet position, and small boxed area indicates an example region used for single-cell tracking in panels D and J. **(B,H)** Change in area of Syn OS-B’ and Chfl MS-1 colonies on day 4 vs day 0. **(C,I)** Change in distance moved towards the light of Syn OS-B’ and Chfl MS-1 colony centers on day 4 vs day 0. Data from 12 biological replicates in 2 experiments. **(D,J)** Microscope images of tracked Syn OS-B’ cells and Chfl MS-1 filaments illuminated from the right, with coloured lines indicating tracks. In panel D, cells were tracked with Trackmate (Fiji) and paths coloured from blue to red over time, while in panel J, the tips of filaments were tracked with custom software (see methods) and each track given a unique color. **(E,K)** Speed of Syn OS-B’ and Chfl MS-1 cells in the dark and light. **(F,L)** Motility bias (proportion of movement towards light) of Syn OS-B’ and Chfl MS-1 cells. Data based on >100 Chfl MS-1 tips or >300 Syn OS-B’ cells. Wilcoxon rank sum test was performed for each panel: ns=P>0.05, **=P<0.01.

Individual Syn OS-B’ cells exhibited “twitching motility” (Figure 1D & Video S3), which is typical of T4P-dependent surface motility in cyanobacteria [33], and moved primarily along their long axis [32]. Chfl MS-1 filaments moved back and forth, parallel to their long axis, often forming bundles (Figure 1J & Video S4). Syn OS-B’ cells and Chfl MS-1 filaments exhibited similar speeds of 0.1-0.15 µm/s that did not differ between the light and dark (Figure 1E&K). Only Syn OS-B’ cells developed a motility bias towards the light (Figure 1F&L). Syn OS-B’ responded to illumination with a small and temporary (∼20 second) increase in speed and a slower (∼2 minute) development of motility bias, and to the dark transition with a temporary (∼30 second) decrease in speed and gradual loss of motility bias (Video S3, Figure S4A&B). Chfl MS-1 speed and motility bias did not change under light transitions (Video S4, Figure S4C&D).

### Interactions between Syn OS-B’ and Chfl MS-1 enhance motility

We next investigated whether interactions between Syn OS-B’ and Chfl MS-1 altered their collective motility. We focused on the role of type IV pili in motility and direct interactions between species in binary consortia, as well as indirect interactions through secreted substances. CYB_2143 in Syn OS-B’ was identified as the homolog of the *Synechocystis PilB1* (slr0063) pilus extension ATPase, and knockout mutants were created using double homologous recombination (Figure S7A&B). These *ΔpilB* mutants still normally expressed genes up and downstream of *pilB* (Figure S8), lacked T4P when imaged by TEM (Figure S7C, S9&10), and exhibited no movement in standard photomotility assays (Figure 2 and S7D).

**Figure 2.**
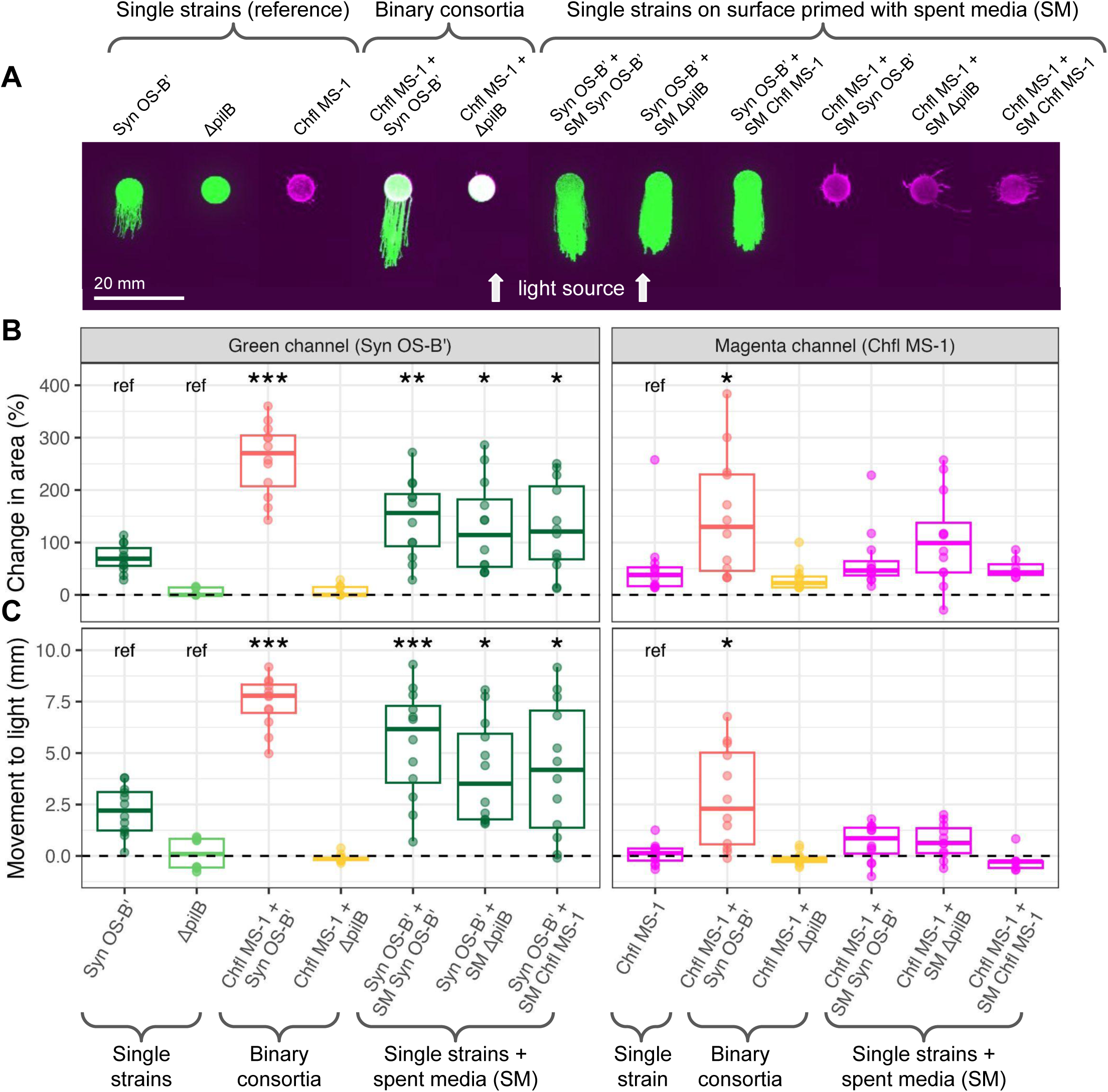
Interactions between Syn OS-B’ and Chfl MS-1 enhance motility. **(A)** Fluorescent scans of collective motility 4 days after exposure to directional (from bottom) white light. From left to right: single strains Syn OS-B’, *ΔpilB* mutant of Syn OS-B’, and Chfl MS-1; binary consortia of Chfl MS-1 mixed with Syn OS-B’ or *ΔpilB* 1:1; and single strains + spent media including Syn OS-B’ or Chfl MS-1 on surfaces primed with filtered spent media (SM) from Syn OS-B’, *ΔpilB*, and Chfl MS-1 cultures. **(B)** Percentage change in the covered colony area of the single strains or binary consortia in panel A after 4 days of light relative to their starting area on day 0. The green (phycocyanin) channel measuring Syn OS-B’ and *ΔpilB* movement is on the left, and the magenta (bacteriochlorophyll) channel measuring Chfl MS-1 is on the right. Colours of dots and boxplots represent different species or consortia, and are labeled at the bottom of panel C. **(C)** Distance moved towards the light (mm) of the center of the colony area of the same consortia as in panel B. Data in panel B and C is from 7-12 biological replicates from 2 separate experiments. Paired student’s t-tests were performed using the single-species with no spent-media (Syn OS-B’, *ΔpilB,* and Chfl MS-1) as the references (ref). The Benjamini-Hochberg adjustment to control false discovery rate was applied per reference group and adjusted p-values reported as follows: *=P<0.05, **=P<0.01, ***=P<0.001.

When *ΔpilB* was mixed in binary consortia with Chfl MS-1, both species moved to the same extent as when alone, suggesting minimal interaction. By contrast, mixing WT Syn OS-B’ with Chfl MS-1 resulted in an increased area covered by each species compared to when they were alone (Figure 2A&B), suggesting a cooperative interaction. Each species also moved further towards the light on average when in the binary consortium (Figure 2C). This resulted in Chfl MS-1 displaying an apparent phototactic bias in the binary consortia. The dynamics of each species in binary consortia initially reflected their movement when alone, with Chfl MS-1 projections emerging in random directions by 6 hours, followed by Syn OS-B’ moving towards the light after 12-24 hours (Video S5, Figure S6). However, after 24 hours, the cooperative effect was more apparent, with Syn OS-B’ and Chfl MS-1 both moving towards the light together. To determine whether secreted substances contributed to the cooperative motility, we collected and filtered spent media from 24-hour single-species cultures and added it to the agarose surfaces before the motility assays. Syn OS-B’ motility was increased by spent media from Syn OS-B’, *ΔpilB*, and Chfl MS-1, both in the area covered and in movement towards the light (Figure 2). By contrast, Chfl MS-1 motility was not significantly affected by spent media.

Our results, therefore, suggest that (i) Syn OS-B’ motility benefits more from some secreted substances than Chfl MS-1, and (ii) direct physical interactions between motile Syn OS-B’ and Chfl MS-1 (including their secretions) result in the greatest cooperative effect.

### Interactions between Syn OS-B’ and Chfl MS-1 create ordered biofilms

To investigate physical interactions between the species, we used fluorescence microscopy to visualize the cellular organization in Syn OS-B’ colonies alone or with Chfl MS-1. Within Syn OS-B’ projections towards the light, individual cells were often well separated and randomly oriented (Figure 3A&D). However, in the projections from binary consortia of Syn OS-B’ and Chfl MS-1, the Syn OS-B’ cells were organized in parallel arrays between Chfl MS-1 filaments (Figure 3B&E, Figure S11). When Syn OS-B’ was replaced in the synthetic consortia by *ΔpilB*, *ΔpilB* cells were also packed in arrays but were less parallel to the light source and we noticed that a few *ΔpilB* cells were transported outside of the starting area (Figure 3C&F, Figure S12).

**Figure 3.**
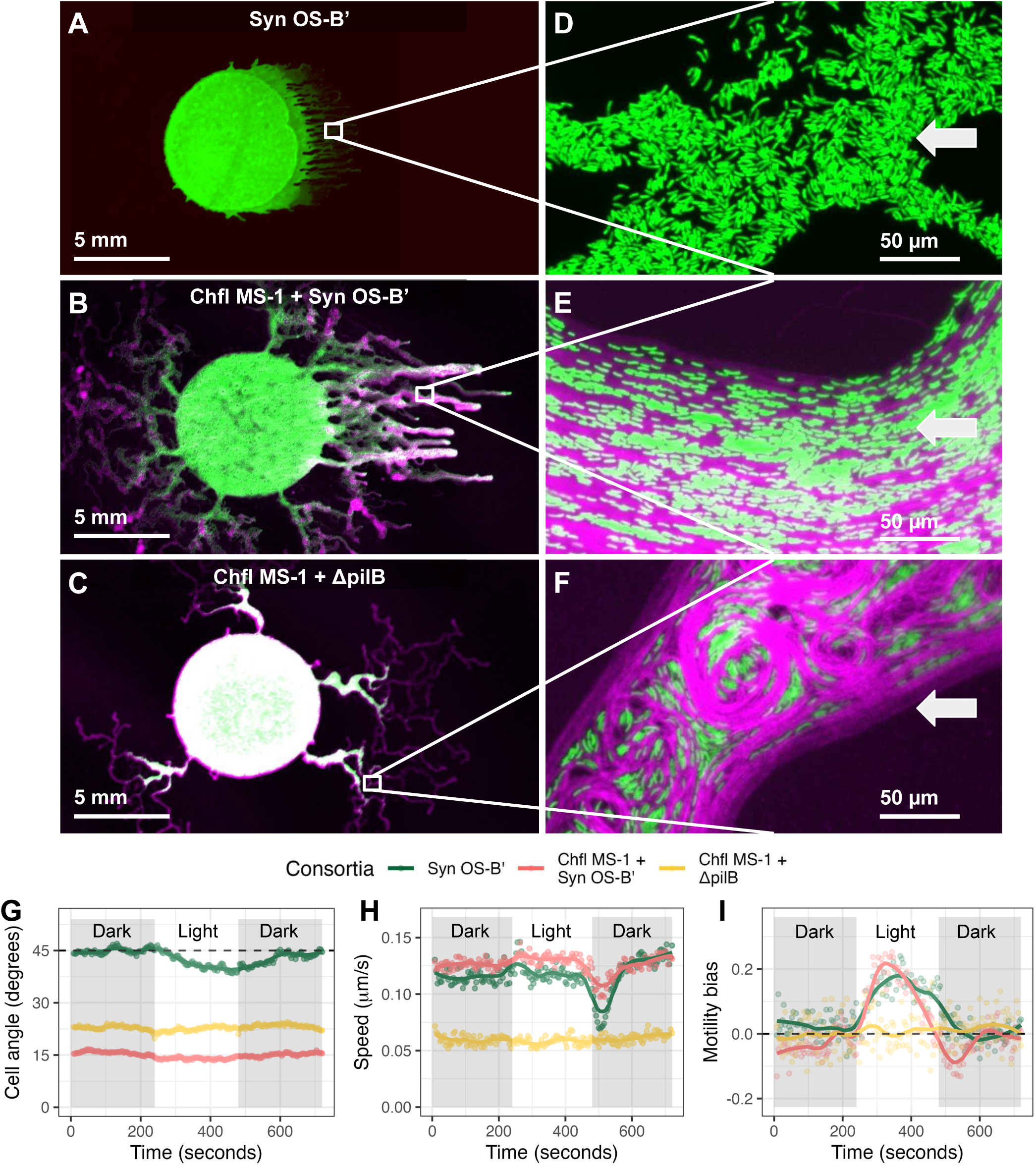
Interactions between Syn OS-B’ and Chfl MS-1 create ordered biofilms. **(A-C)** Fluorescence scans of Syn OS-B’ (top), Chfl MS-1 + Syn OS-B’ (middle), and Chfl MS-1 + *ΔpilB* (bottom) after two days of incubation with white light from the right. Green regions = Syn OS-B’ or *ΔpilB*, magenta regions = Chfl MS-1, white regions = both species. **(D-F)** Fluorescence microscope images of the indicated regions in panels A-C showing Syn OS-B’ or *ΔpilB* rods in green (ex. 540-580 nm / em. 608-682 nm) and Chfl MS-1 filaments in magenta (ex.450-490 nm / em. >515 nm). **(G-J)** Dynamics of Syn OS-B’ and *ΔpilB* movement during light transitions (gray regions=dark, white region=directional illumination with white light). Green = Syn OS-B’, Pink = binary consortium of Chfl MS-1 + Syn OS-B’, Yellow = binary consortium of Chfl MS-1 + *ΔpilB*. **(G)** Average (mean) angle of the long axis of Syn OS-B’ or *ΔpilB* cells relative to the light direction (0° = parallel, 45° = random, 90° = perpendicular). **(H)** Mean speed of Syn OS-B’ or *ΔpilB* cells over time. **(I)** Motility bias of Syn OS-B’ or *ΔpilB* cells over time. Data from 10 biological replicates in 4 experiments with >300 total cells tracked at each time point per condition.

The strong alignment of Syn OS-B’ cells with the light direction in binary consortia suggests an interaction between motile Syn OS-B’ and Chfl MS-1 cells.

Next, we investigated the consortia under light transitions. After transferring the consortium to the dark, Chfl MS-1 filaments were observed at the tip of the projection and moved around the Syn OS-B’ cells (Video S6). However, after transitioning to the light, Syn OS-B’ cells moved toward the light between Chfl MS-1 filaments, accumulated near the tip, and appeared to drive the projection toward the light. *ΔpilB* cells in binary consortia showed minimal movement in the dark or light (Video S7). Syn OS-B’ cells at the tip of the projection were too densely packed to allow reliable cell tracking. However, in the middle of the positive phototactic projections, we could track Syn OS-B’ using manually trained and automated image segmentation in ImageJ. As before, we preceded the cell tracking with a 10-minute dark incubation, after which Syn OS- B’ cells in the phototactic projections were randomly oriented (mean angle of ∼45° from the prior light source). Syn OS-B’ cells in the consortium with Chfl MS-1 were more parallel to the light (∼15°) (Figure 3G). *ΔpilB* cells within binary consortia that had moved towards the light were selected for comparison, and showed only moderate light alignment of ∼25° (Figure 3G). Upon directional illumination, Syn OS-B’ cells reoriented slightly toward the light (from 45° to 40°) and returned to a random orientation (45°) in the dark. Syn OS-B’ and *ΔpilB* cells in binary consortia did not appear to reorient in the light, perhaps due to restrictions by Chfl MS-1 filaments (Figure 3G & Videos S8&9). Syn OS-B’ cell speed was similar when alone or in binary consortia, but significantly higher than *ΔpilB* in consortia (Figure 3H). Some movement of *ΔpilB* cells was observed in binary consortia but this was likely due to the movement of neighboring Chfl MS-1 filaments. During the transition from light to dark conditions there was a noticeable but temporary decrease in Syn OS-B’ speed, as previously observed [32], which was mitigated in binary consortia with Chfl MS-1, and not exhibited by *ΔpilB* (Figure 3H). As expected, Syn OS- B’ increased its motility bias over the first 60-120 seconds of illumination when alone and in the binary consortium (Figure 3I), however, this bias declined after 120 seconds in both cases. The decline in Syn OS-B’ bias was particularly striking in binary consortia suggesting that Chfl MS-1 may initially restrict Syn OS-B’ movement. However, this is likely temporary as the collective motility assays (Figure S6B) and longer videos (Video S6) indicated that Syn OS-B’ continued to move steadily towards the light over longer periods.

### Interactions between Syn OS-B’ and Chfl MS-1 create stronger biofilms

To observe biofilm formation under liquid, we incubated Syn OS-B’, *ΔpilB*, and Chfl MS-1 alone and in binary consortia in DH10 + 250 mg·L^−1^ tryptone under constant white light in 384-well plate format. Imaging the bottom of the wells after 8 days (Figure 4A) showed various patterns. Syn OS-B’ and *ΔpilB* produced a diffuse distribution pattern while Chfl MS-1 produced a reticulate network composed of bundles of filaments. In binary consortia, Syn OS-B’ formed clusters within a Chfl MS-1 network, and *ΔpilB* + Chfl MS-1 binary consortia exhibited aggregation and colocalization of the two species. Next, we measured total culture density, density of the planktonic cells (resuspended by gentle pipetting), and biofilm-associated cells by crystal violet assay in 96-well tissue-culture treated plates. The total culture densities of the binary consortia and single-species Syn OS-B’ and *ΔpilB* cultures were similar, but the Chfl MS- 1 culture density was significantly lower (Figure 4B). The planktonic cell densities of the Syn OS-B’ and *ΔpilB* cultures were significantly higher than Chfl MS-1 or the binary consortia, suggesting that Syn OS-B’ and *ΔpilB* did not attach to the plate surface as well as Chfl MS-1, and that the presence of Chfl MS-1 improved their retention in the biofilm (Figure 4B). Indeed, in the consortia, the biofilm levels were significantly higher than any of the single-species cultures (Figure 4B). We also measured the dynamics of biofilm formation over four days. We found detectable biofilm levels after 6 hours (Figure S13), with maximal biofilm development occurring between 6-24 hours, concurrent with a decrease in the planktonic fraction.

**Figure 4.**
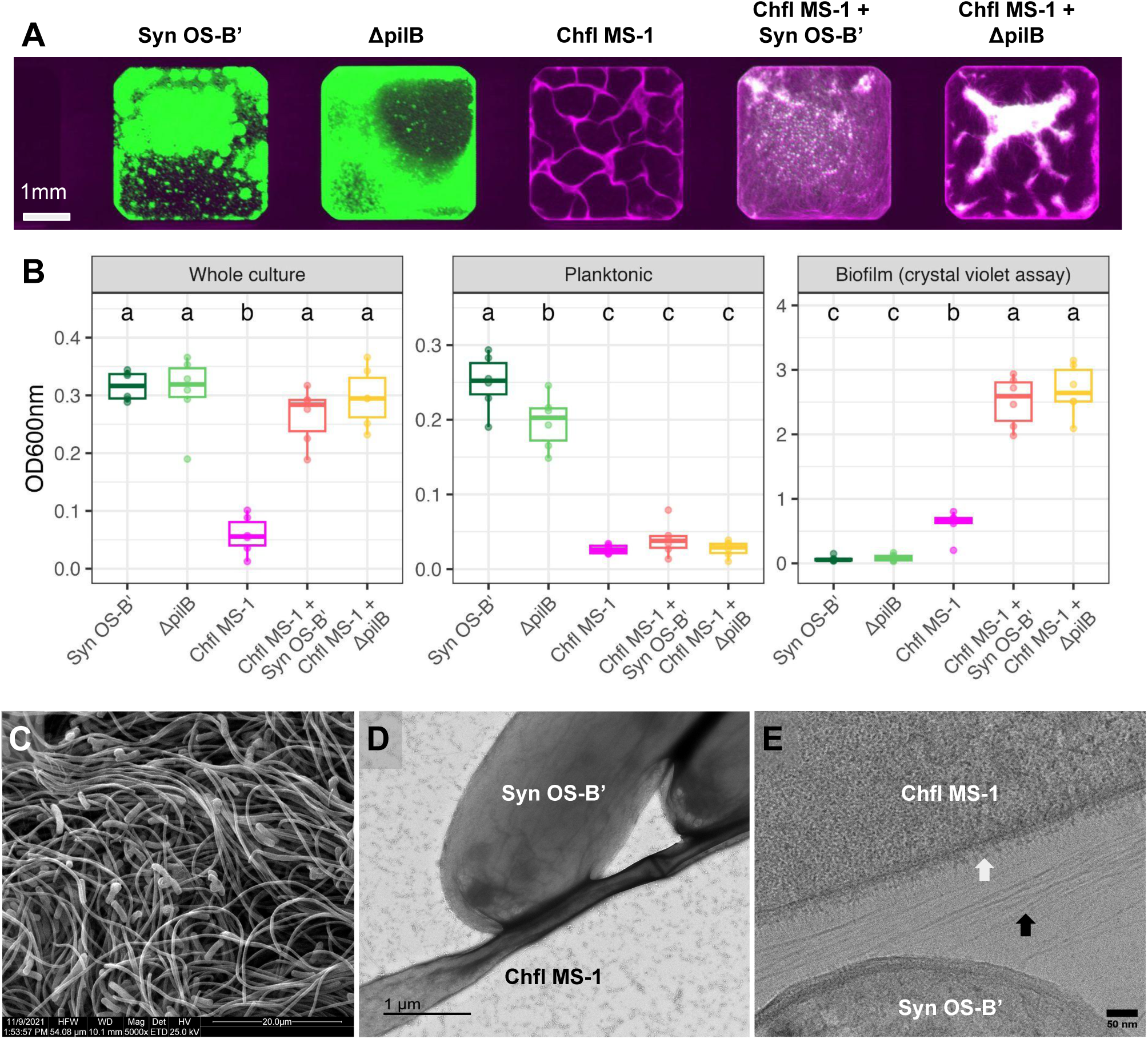
Interactions between Syn OS-B’ and Chfl MS-1 create stronger biofilms. **(A)** Representative scanning fluorescence images of the bottom of culture wells in a multiwell plate after 8 days growth in DH10 + 250 mg/L tryptone. Green regions indicate Syn OS-B’ and *ΔpilB* (ex. 658 nm / em. 710 nm), magenta regions indicate Chfl MS-1 (ex. 784 nm / em. 832 nm), white regions indicate both species. **(B)** Boxplot and individual measurements of OD600nm values from various fractions of the cultures: whole culture (left), planktonic fraction (middle), biofilm fraction after crystal violet assay (right). Data from 6 biological replicates in one experiment with additional data in Figure S13. Compact letter display (letters at top of plot) indicates statistical differences at *P*<0.05 in group means after Tukey’s HSD test. **(C)** Syn OS-B’ + Chfl MS-1 consortia imaged by SEM at 5000x. **(D)** Syn OS-B’ + Chfl MS-1 consortia stained with uranyl-acetate and imaged by TEM at 30,000x. **(E)** Syn OS-B’ + Chfl MS-1 consortia plunge-frozen in liquid ethane, milled by cryogenic focused-ion beam SEM (cryoFIB-SEM), and then imaged by cryo-EM. White arrow indicates a fuzzy external layer (EL) to the cell membrane of Chfl MS-1. Black arrow indicates long, thin, pili-like filaments (6-8 nm diameter).

To visualize physical interactions within the biofilm, we prepared binary consortia of Syn OS-B’ and Chfl MS-1 in liquid culture for scanning (SEM), transmission electron microscopy (TEM), and cryo-electron microscopy (cryo-EM). SEM images revealed a close association between Syn OS-B’ and Chfl MS-1, with Chfl MS-1 filaments forming the bulk of the biofilm (Figure 4C & S14). TEM images of the associations showed increased electron density between the cells.

This might be interpreted as secreted EPS between species but needs further experimentation (Figure 4D & S15) [34]. Cryo-EM revealed a fuzzy external layer on the surface of Chfl MS-1 cells and many pilus-like structures between Syn OS-B’ and Chfl MS-1 (Figure 4E). Together, these results suggest that Chfl MS-1 and Syn OS-B’ establish tight interactions producing a more robust biofilm when together.

## Discussion

We investigated how bacterial motility responses to light can shape the architecture of microbial mats using two dominant members of Yellowstone hot spring mat communities. We found comparable behavior of Syn OS-B’ and Chfl MS-1 to other observed Cyanobacteria and Chloroflexota. By studying the motility and arrangement of these species in binary consortia, we revealed some unique emergent properties. These insights could enhance our understanding of the dynamics of natural microbial mat architecture, and help to develop synthetic consortia for uses in industry.

## Syn OS-B’ and Chfl MS-1 behavior is comparable to other Cyanobacteria and Chloroflexota

Chfl MS-1 exhibited surface motility at speeds consistent with other hot spring Chloroflexota [35], and formed gliding aggregates of filaments similar to *Chloroflexus aggregans* [36]. Chfl MS-1 did not exhibit phototaxis, which, to our knowledge, has not been reported in the Chloroflexota, however, light does increase *C. aggregans* aggregation rate [37]. The genes responsible for Chfl MS-1 motility were not investigated here, but the Chfl MS-1 genome [30] does contain TadE/G genes of the tight adherence pilus family, which is involved in the motility of *Liberibacter crescens* [38].

As we reported previously [32], Syn OS-B’ was similar in speed and phototactic bias to other characterized cyanobacteria, particularly those from Yellowstone hot springs [27]. Here, we found that disruption of the homolog of *pilB* in Syn OS-B’ (CYB_2143), resulted in loss of T4P and motility, as found in *Synechocystis* sp. PCC 6803 [39]. Syn OS-B’ did not form adherent biofilms, as observed in laboratory strains of model cyanobacteria such as *Synechocystis* PCC 6803 and *Synechococcus elongatus* PCC 7942 grown under normal conditions, but differing from some related wild isolates, which do form biofilms [40], [41]. The mutant lacking *pilB* was slightly less abundant in the planktonic phase than Syn OS-B’, suggesting that pili might prevent cell sinking, as observed in marine cyanobacteria [42], or inhibit biofilm formation, as found in *S. elongatus* [43].

## A unique synthetic system for studying emergent biofilm properties

This binary consortium has relatively unique features useful for mixed-species biofilm research, including: (i) strains recently isolated from well-studied microbial mats, with few generations to acquire laboratory mutations; (ii) a member (Syn OS-B’) that is naturally transformable allowing mutant generation to test gene function e.g. *ΔpilB*; (iii) two members with different morphologies (rod-shaped vs filamentous), motilities (phototactic twitching vs gliding), and fluorescent pigments (allowing species to be distinguished); and (iv) emergent or cooperative properties in both surface motility and biofilm formation when the species are combined.

Cooperative microbial motility often depends on the release of surfactants, such as rhamnolipids in *P. aeruginosa* [44]. We found that Syn OS-B’ motility was slightly improved by supplementing the agar surface with spent medium from Syn OS-B’, *ΔpilB*, and Chfl MS-1 suggesting Syn OS- B’ surfactant production may limit its surface motility. Indeed, the finger-like collective projections we observed for Syn OS-B’ can indicate collective movement limited by surfactant production [45]. Further investigation of the chemical nature of these surfactants could help elucidate their role in cross-species cooperation. In addition, direct physical interactions between cells also appear crucial to the cooperative effect. This might partly be explained by the branching-exploratory behavior of Chfl MS-1, which can maximize colonization [46], combined with the phototactic directionality of Syn OS-B’. Importantly, Chfl MS-1 exhibited the emergent property of positive bias towards the light only when mixed with Syn OS-B’. Mathematical models that incorporate morphological information, directional bias, and motility-promoting substances could be useful for understanding this cooperative phenomenon.

Mixed-species biofilms are often more disordered than single-species biofilms and less robust, for example to predation or invasion [47], [48]. By contrast, we found that Syn OS-B’ was more ordered and aligned with the light source when mixed with Chfl MS-1 than when alone. This may result from repeated collisions between Chfl MS-1 filaments and the phototactic Syn OS-B’ rods leading to the alignment in the primary direction of movement. This long-range ordering via collisions has been observed in biofilms of filamentous *E. coli* [49], and mathematical models based on filamentous cyanobacteria [50]. Statistical models of biofilm architecture have found that the cellular length:width ratio is amongst the most important determinants of biofilm structure [51], corroborating the finding that filamentous Chfl MS-1 had a large impact on Syn OS-B’ arrangement.

In addition to increased motility, we observed higher biofilm formation in the binary consortia than in single-species cultures. Increased biofilm formation in species mixtures has been found in bacteria isolated from the soil and human gut [52], [53]. In our case, this is at least partly due to stronger aggregation and substrate attachment in binary consortia, rather than just increased growth, as Syn OS-B’ and *ΔpilB* decline in the planktonic fraction when mixed with Chfl MS-1. Chfl MS-1 formed biofilms when alone, suggesting it is the driver of biofilm adherence. This agrees with other findings that filamentous cells often form the biofilm core or help stabilize its structure [54].

## Cyanobacteria-Chloroflexota biofilms in natural and industrial settings

In hot springs, Chloroflexota are reported to mainly acquire organic carbon from cyanobacteria, often as excreted organic acids [55], [56]. Moving and associating with cyanobacteria may therefore provide an advantage for Chloroflexota. Indeed, vertically oriented Chloroflexota filaments have been observed amongst cyanobacterial cells in hot spring mats, particularly during mat regrowth [17], potentially suggesting alignment by phototactic cyanobacteria moving towards or away from the light. Alternatively, vertical migrations of Chloroflexota in Octopus spring mats might also be due to positive aerotaxis [17], which has been characterized in *Chloroflexus aggregans* in the laboratory [57]. We performed motility experiments on soft agarose surfaces exposed to air in part to avoid the potential confounding effect of oxygen gradients. However, future studies could recapitulate three-dimensional biofilm architecture in microfluidic devices with flow, and extend our photomotility experiments to more complex and realistic environments.

Chloroflexota are considered the main contributors to mat structure in several Yellowstone hot springs because they are abundant in structures above the mat surface such as ‘nodules’ or ‘streamers’, and continue to form a mat when cyanobacteria are diminished [17]. This is consistent with our results that Chfl MS-1 can form strong biofilms without Syn OS-B’ but not vice versa. However, Boomer *et al*. found that *Synechococcus* and *Thermus* were the first genera to form biofilms on glass rods suspended above hot spring mats, with Chloroflexota and filamentous cyanobacteria only contributing to the biofilm months later [58]. Future work could therefore test the motility and biofilm formation of other mat members, either in isolation or in mixed-species biofilms. It is also important to note that many microbial mats are dominated by filamentous cyanobacteria [18], [59], which may form adherent biofilms and long-range order without interacting with other filamentous species.

Chloroflexota and Cyanobacteria don’t only associate in natural habitats. Zhang *et al*. found they were the main contributors to nutrient removal in municipal wastewater treatment photobioreactor experiments [60]. These phyla have each received considerable attention in wastewater treatment [61], [62], and it has been suggested that their combination could provide advantages of reduced greenhouse gas emissions and better surface adherence or settleability to reduce biomass separation costs [63], [64]. Cyanobacterial biofilm formation also offers advantages in increasing bioenergy and biofuel productivity [65], and the addition of heterotrophic strains can increase cyanobacterial attachment [66]. Our results that binary consortia of cyanobacteria and Chloroflexota isolates cooperate to form robust biofilms may therefore inform phototrophic biofilm cultivation strategies for commercial applications.

## Conclusion

We found that Cyanobacteria and Chloroflexota isolates that co-occur in hot springs show different motility responses to light but can cooperate to form robust biofilms and migrate over larger distances. These results provide insights into physical interactions within hot spring mats and highlight understudied features of biofilm communities, including directional movement via phototaxis, the emergence of long-range cellular order, and cooperation in both motility and biofilm formation.

## Methods

### Strains and culture conditions

*Synechococcus* sp. JA-2-3B′a(2-13) (OS-B′) was originally isolated by filter cultivation from a 51-61°C Octopus Spring mat core sample [67]. *Chloroflexus* sp. MS-CIW-1 was isolated from a 60°C Mushroom Spring mat core sample by serially picking colonies and streaking them on PE medium (0.4% agarose) plates. Both strains were stored at −70°C in 20% (wt/vol) glycerol stocks prior to culturing. Syn OS-B’ cultures were maintained in DH10M medium (medium D [68] plus 10 mM HEPES adjusted to pH 8.2, with added 250 mg·L^−1^ bacto-tryptone, and 750 mg·L^−1^ sodium bicarbonate). Chfl MS-1 was maintained in PE medium [69]. Both species were incubated at 50°C under continuous light at 50 μmol·m^−2^·s^−1^ (measured with a LI-COR [model Li-189] radiometer) in an Algaetron ag130 incubator (Photon Systems Instruments) or similar incubators. To determine axenicity, we plated the cultures, incubated them on DH10M agarose plates with added yeast extract (100 mg·L^-1^) and checked for any non-Syn OS-B’ or non-Chfl MS-1 colonies.

### Collective motility assay

Cultures in late-log phase (OD600nm of Syn OS-B’ and Chfl MS-1 of ∼0.6 and ∼0.3, respectively) were concentrated to an OD600nm of 2.0 using a spectrophotometer (Tecan M1000 Pro), centrifugation, and resuspension in DH10 (medium D [68] plus 10 mM HEPES adjusted to pH 8.2). 5 µL of these concentrated cultures were plated on DH10 solidified with 0.4% (wt/vol) agarose in bacteriological petri dishes with tightly fitting lids (Falcon 25369-022) and allowed to dry for ∼10 min in a laminar flow cabinet. Plates were then inverted and incubated at 50°C with unidirectional daylight white LED illumination (spectrum in Figure S3) at 60 μmol·m^−2^·s^−1^. Plates were imaged using a fluorescent scanner (Azure biosystems) at 0, 6, 24, 48, 72, and 96 hours after incubation. Two fluorescent channels were used: excitation at 658 nm or 784 nm with emission capture at 710 nm or 832 nm with 40 nm bandpass for Syn OS-B’ or Chfl MS-1, respectively. Images were processed using custom ImageJ macros to identify the regions where Syn OS-B’ and Chfl MS-1 had moved. The experimental setup metadata file describing where drops were placed on the plates and this extracted colony distribution data were matched using the Hungarian algorithm for solving the linear sum assignment problem in R version 4.3.1.

### Single-cell time-lapse video microscopy

Single-cell motility measurements were captured by Time-lapse video microscopy (TLVM). The temperature was maintained at 50°C using a custom environmental chamber (HaisonTech), and illumination was provided by rows of white LEDs (spectra in Figure S1) positioned 10 cm from and at 5° above the agarose surface. The light intensity measured at the cells’ position was 60 μmol·m^−2^·s^−1^. Cells were prepared as described above for collective phototaxis measurements with Chfl MS-1 filaments tracked from a lower density droplet within hours after plating, while Syn OS-B’ cells were tracked after 2-4 days and amongst cells that had already displayed positive phototaxis. All petri dishes remained closed and inverted during recording with a minimum 10-minute incubation at 50°C in darkness prior to the 4- to 12-minute recordings at ×200 magnification and 10-15 frames per minute. All videos were made using a Nikon Eclipse TE300 inverted microscope attached to a Coolsnap Pro monochrome camera (Media Cybernetics, Silver Spring, MD, USA), and images were collected by MetaMorph software (v4.6r5 and 6.2r6; Molecular Devices). Example videos can be found at https://www.youtube.com/channel/UCX0gm-79tZplREzHteapEWA.

### Syn OS-B’ tracking

Image stacks were preprocessed using Image J v1.53f51 [70] by subtracting the background with a rolling ball radius of 4. Cells were tracked using the ImageJ plugin Trackmate [71] with a thresholding filter applied and filtering to blobs of >25 pixels. The Kalman tracker was selected with the initial search radius set to 15, the subsequent search radii to 15, and the maximum frame gap to 3. The “spots” file was exported to csv and imported into R version 4.3.1 for preprocessing and analysis [72]. Speed was calculated as distance (in μm)/time (in s). Motility bias was calculated as parallel displacement (in μm)/distance (in μm).

### Chfl MS-1 tracking

We developed a custom tracking algorithm for filamentous bacterial tracking in LabView programming language, with a convenient user interface. The timelapse 2D images were corrected by pixelization, integrated FFT noise filtering, simple Kalman filtering, adjacent pixel averaging and 4th-degree polynomial curve correction in X direction and adaptive pixel intensity inversion to ensure that bacterial intensity is above the background. The tracking was performed on the original frame on 10 filaments per video. The filament ends were picked manually by positioning the cursor in an interactive image and after thresholding, the algorithm tracked the filament tip between frames. The end pixels were found by the “grassfire” method and their center of mass was taken as the true end. The tracks were plotted in real time with a set delay, to view the tracking quality in real time and ensure fidelity. After the last frame, the tracks were saved in a text file and analyzed in R version 4.3.1 to calculate speed and motility bias as above.

### Biofilm quantification

Biofilm formation was quantified using the crystal violet assay [73], with modifications as in [74]. Briefly, Syn OS-B’, *ΔpilB*, Chfl MS-1, and binary consortia were inoculated at OD600nm of 0.4 into DH10 or DH10 + BT media in 96-well polystyrene plates and incubated under continuous white light at 50 μmol·m^−2^·s^−1^ and 50°C. After 0-4 days of incubation, OD600nm absorption measurements were taken on the total culture. Then, cultures were briefly disrupted by pipetting, and 80% of the culture volume was transferred to a new plate. OD600nm was measured in the 20% of the remaining volume and called “Non-planktonic OD600nm”, and in the transferred volume, called “Planktonic OD600nm”. Plates were washed three times by adding 200 μl of Milli-Q water, pipetting up and down, then removing and discarding 200 μl. Then, 200 μl of aqueous 0.1% crystal violet was added to each well and the plate incubated at room temperature for 20 minutes. Plates were washed 3 times with Milli-Q water as before, then 200 μl of 95% ethanol was added to each well and the plate agitated for 5 minutes. OD600nm was then recorded and reported as the Biofilm (CV) level.

### Scanning electron microscopy

A 5 µl aliquot of a late log phase culture of Syn OS-B’ + Chfl MS-1 was added to the surface of an SEM stub (Ted Pella), and a Kimwipe tissue (Kimtech) was used to blot away excess liquid. The SEM stub was then immersed in 100% methanol for 5 minutes, rinsed with 100% ethanol, and immersed in ethanol for 10 minutes. The SEM stubs then underwent critical point drying with LCO2 in DCP-1 (Denton Vacuum) in 200-proof ethanol at a maximum of 50-60℃ and 9-10 MPa, with depressurization at 300-700 kPa per minute. Gold sputtering from pure Au target was performed in Denton Desk IV (Denton Vacuum) under argon gas at a pressure of 6 kPa for 200 seconds and a low current of 7 mA. The samples were imaged in Quanta 200 (FEI) SEM with 25 kV accelerating voltage in high vacuum with secondary emission detector.

### Transmission electron microscopy

300 mesh Formvar and carbon-coated copper grids (FCF300-Cu, Ted Pella) were irradiated with low-energy Al plasma glow to create a hydrophilic surface in Denton Desk II instrument (Denton Vacuum). The Syn OS-B’ + Chfl MS-1 binary consortium was gently mixed and 5 µl drops were deposited on the shiny side of the grids and left for 3 minutes before negative staining was performed using standard methods. This involved dipping the grid face-down consecutively onto two DI H2O droplets, followed by rinsing in 3 drops of 1% uranyl acetate, and resting to stain for 1 minute before blotting away the remaining uranyl acetate and leaving the sample to dry completely. The grids with bacteria were imaged in JEOL JEM-1400 with 120 kV accelerating voltage equipped with a Gatan Inc. OneView 4kX4k sCMOS camera.

### Cryo-electron microscopy

A binary consortium of Chfl MS-1 and Syn OS-B’ was grown for 6 days in DH10 medium in a sealed 96 well plate at 50°C and under 12:12hr light-dark cycles with LED daylight white illumination at 60 μmol·m^−2^·s^−1^. After 11 hrs of the dark period, the biofilm was resuspended gently by pipetting, diluted 9-fold in DH10, and 3 μl of cell suspension transferred onto 200 mesh Carbon Quantifoil R1.2 Gold TEM grids, which were glow-discharged using a Pelco EasiGlow to enhance hydrophilicity. Cells were allowed to settle for 30 seconds, followed by single-sided blotting for 4-8 seconds and vitrification in liquid ethane using a Leica EM GP2 robotic plunge-freezer. Samples were transferred cryogenically into a Thermo Fisher Aquilos2 Dual Beam cryogenic focused ion beam scanning electron microscope (cryo-FIB/SEM) to generate 200 nm thin lamellae. The stage temperature was -185 °C to maintain vitreous ice during milling. Grids were sputter-coated with platinum for 25 seconds, followed by a protective layer of organo-platinum for 2 minutes using the integrated gas injection system. Milling was performed with the Gallium-ion beam at 30 kV and currents decreasing from 500 to 13 pA until 500 nm thickness, then at 10 pA to 200-300 nm. SEM inspection of the lamellae and final polishing was conducted with a 2 kV and 13-25 pA electron beam. Samples were transferred to a Thermo Fisher Titan Krios G3i cryo-TEM and screening images were collected using SerialEM 4.11.12 on a K3 direct electron detector (Gatan Inc.) with a Bioquantum energy filter using 20 eV slit width. Images were collected at 42,000x nominal magnification with 2.6 Å/pix, an exposure of 0.3 seconds, and a defocus of -3 µm. Images were binned 4x for visualization, and ultrastructure measurements were performed using IMOD 4.11.12 [75] or FIJI [70].

## Acknowledgments

F.B. acknowledges support from the University of Chicago, Carnegie Institution for Science, and a BBSRC-NSF/BIO collaborative research grant (award number 1921429). V.C. was supported by a grant from the National Aeronautics and Space Administration (80NSSC19K0462). C.R., D.B., and A.G. acknowledge support from NSF/BIO 1921429 and from the Carnegie Institution for Science. This research was partially supported by the DOE Office of Biological and Environmental Research, Biological Systems Science Division, FWP 74915, and by NIH S10 Award Number 1S10OD028536-01 from the Office of Research Infrastructure Programs (ORIP). Part of this research was performed on a project award (10.46936/expl.proj.2021.60171/60008194) from the Environmental Molecular Sciences Laboratory, a DOE Office of Science User Facility sponsored by the Biological and Environmental Research program under Contract No. DE-AC05-76RL01830. Some of this work was performed at the Stanford-SLAC Specimen Preparation Center (SCSC), which is supported by the National Institutes of Health Common Fund’s Transformative High-Resolution Cryoelectron Microscopy Program (U24GM139166). We thank the Yellowstone National Park Service and David Ward for sampling permit #YELL-5494 awarded to David Ward and Devaki Bhaya, which allowed the collection of mat samples from which Chfl MS-1 and Syn OS-B’ were isolated. We also thank Stanford CSIF and Carnegie Advanced Imaging Facility for access to their TEM and SEM instruments.

## Author contributions

F.B. and D.B. conceived and designed the research. F.B., C.R., V.C., and A.M. carried out experiments. A.M., L.M.J, A.D.P, and J.E.M. provided tools, sample processing, and performed microscopy analysis on samples. F.B. and D.B. analyzed the results and wrote and edited the manuscript with input from all authors.

## Data availability statement

All data presented in this manuscript are available in the supplementary data files along with an R script to reproduce all plots and statistical tests.

## Supplementary methods

### Plasmid construction for *ΔpilB* knockout in Syn OS-B’

A construct containing a thermostable kanamycin resistance cassette (KmTe^R^), flanked by up- and down-stream regions homologous to the *pilB* (CYB_2143) deletion site was built in pGEM®- T easy (Promega). The pT7BT/P cpcC::KmTe plasmid containing the thermostable kanamycin resistance cassette from *Thermosynechococcus elongatus* (KmTe^R^) was a generous gift from Kiyoshi Onai (Nagoya University) (1). The fragments were amplified by PCR with primers 1-3 (Table S1) using Phusion High Fidelity PCR Kit (New England Biolabs, USA) and assembled with NEBuilder® HiFi DNA Assembly Master Mix (USA). The construct was cloned in *E. coli* DH5-α and confirmed by Sanger sequencing through ELIM biopharmaceuticals Inc. (USA). In OS-B’, the *pilB* gene may be included in an operon that includes putative pilus-related genes *pilT* and *pilC* downstream of *pilB*. To avoid a potential polar effect by the disruption of the *pilB* gene, several factors were considered as outlined in (2): i) the KanR was inserted in the same orientation as the genes in the putative operon, ii) no transcriptional terminator in the KanR was included, iii) the KanR was inserted by the 3’ end of the gene but maintaining at least a few codons of the 3’ end of the ORF (101 bp), and iv) the size of the deletion (1,210 bp) is similar to the insertion (878 bp).

### Transformation of Syn OS-B’

Syn OS-B’ cells were grown under continuous light at 50°C on DH10 medium supplemented with 250 mg/l of bacto-tryptone (DH10+BT) (3) until late log phase (OD600nm∼0.6). Cells were adjusted to OD730nm = 60 in 80 µl and incubated with 20 ng/µl of pDNA overnight for natural transformation (1). After incubation, the cells were spread onto two-layer DH10+BT tight-fit bacteriological plates (1% and 0.4% agarose bottom and top layers), and incubated at 50°C under continuous light. After 8-10 days, a kanamycin concentration gradient was created by lifting the bottom agarose layer and adding 40 µL of 30 µg/ml of kanamycin. After 6-8 weeks, antibiotic-resistant colonies were picked and segregated in DH10+BT liquid medium with increasing concentrations of antibiotic to reach homoplasmy, transferring to fresh medium weekly for 3 weeks. The clones were genotyped by PCR using primers 12-14, which bind up and downstream of the homology regions used for recombination, and confirmed by Sanger sequencing. RT-qPCR using primers 4-11 (Table S1) on RNA from Syn OS-B’ and *ΔpilB* cells grown to late-log phase in DH10 medium indicated that there was no obvious polar effect on *pilC*, which is downstream of *pilB*, since it was expressed at similar levels in Syn OS-B’ and *ΔpilB* (Figure S4). Six clones were confirmed and used for motility assays.

### RT-qPCR

Syn OS-B’ and *ΔpilB* were grown in 15 ml of DH10 medium until late log phase (OD600nm∼0.6). Flasks were swirled in liquid nitrogen to chill cultures to <4℃ in <30 seconds, transferred to prechilled 15 ml tubes, centrifuged at 4,000 *g* for 5 minutes, supernatant discarded, and loose pellet transferred to a 1.5 ml tube for a 2-minute 4,000 *g* centrifugation, supernatant removal, and snap-freezing in liquid nitrogen. RNA was extracted using a Quick-

RNA kit (Zymo Cat#R2014) according to the manufacturer’s instructions, with an additional DNAse digestion step with an RNAse-free DNAse set (Qiagen Cat#79254) according to the manufacturer’s instructions. 400 ng of RNA was reverse-transcribed to cDNA using the iScript Reverse Transcription Supermix (Bio-Rad Cat#1708841). cDNA was diluted 10-fold and 4 μL was added to a master mix of 5μL 2× SensiFAST SYBR No-ROX kit reaction mix (Meridian Bioscience Cat#98050) plus 0.5 μL of each 10 mM primer stock (Table S1). PCR amplification was performed over 45 cycles of 95°C for 10 s and 60°C for 30 s with fluorescence quantification at the end of each cycle, followed by melting curve analysis by ramping the temperature from 65°C to 97°C at 0.11°C per second. Absence of gDNA contamination was confirmed by also supplying no-reverse transcriptase cDNA synthesis controls as qPCR templates and finding threshold cycle values >40. qPCR target specificity was confirmed by a single peak in the melt profile with a melt temperature <2℃ from the median for that target. The mean threshold cycle of 4 genes (cysE, La, U32, SufS) served as the housekeeping controls and relative transcript abundances calculated according to the ΔCt method (2^−(Ct target gene − Ct HK mean^)).

---

**Figure S1.**
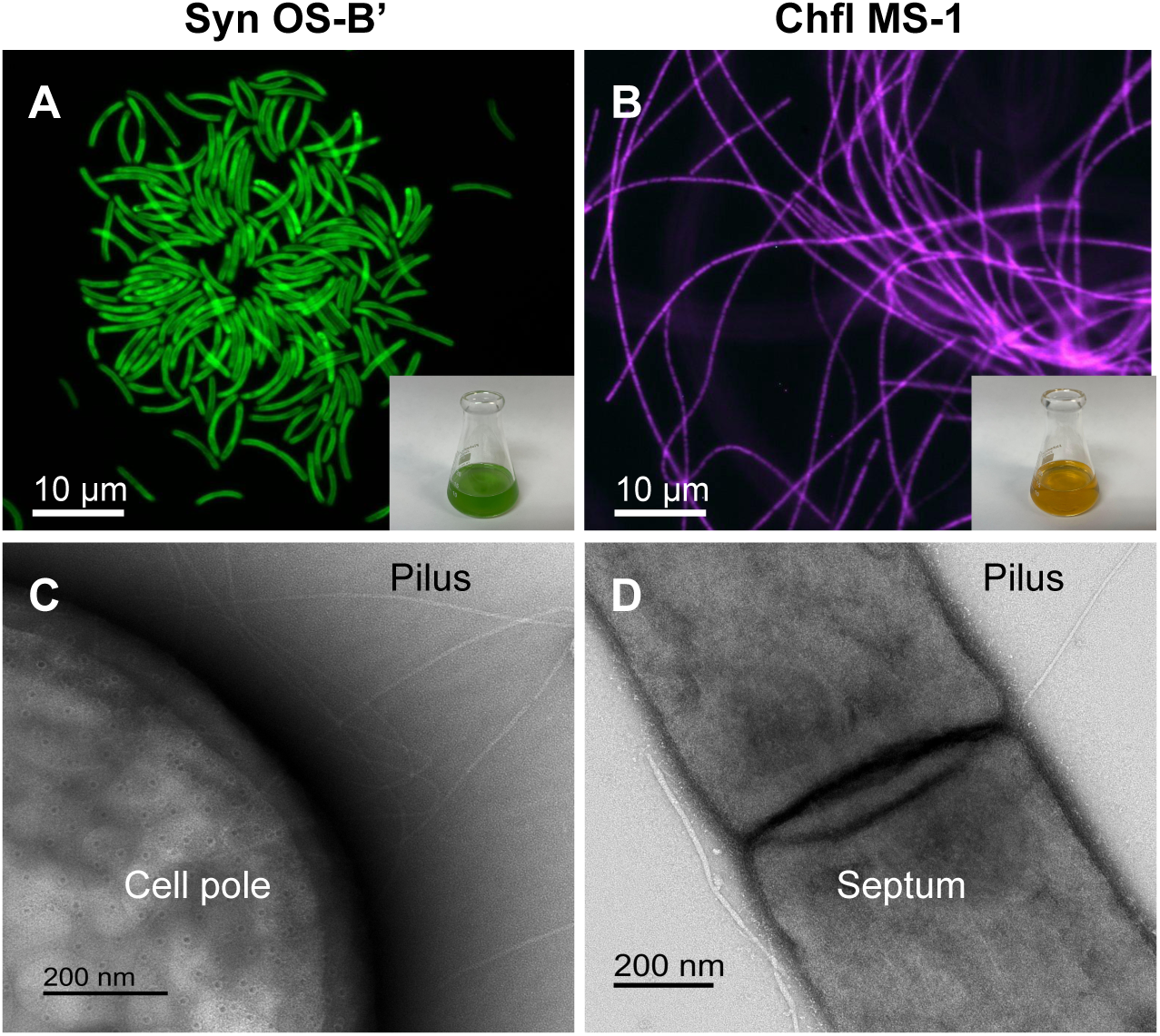
Hot spring isolates Syn OS-B’ and Chfl MS-1 are pigmented and piliated. **(A)** Fluorescence micrograph of Syn OS-B’ (ex. 540-580 nm / em. 608-682 nm); inset: culture of Syn OS-B’ in DH10M medium. **(B)** Fluorescence micrograph of Chfl MS-1 (ex.450-490 nm / em. >515 nm); inset: culture of Chfl MS-1 in PE medium. **(C)** TEM image of uranyl-acetate-stained cell pole of Syn OS-B’ exhibiting T4P (8 nm diameter). (**D**) TEM image of Chfl MS-1 showing the septum between adjacent cells in a filament and a pilus (7 nm diameter) emerging from near the septum.

**Figure S2.**
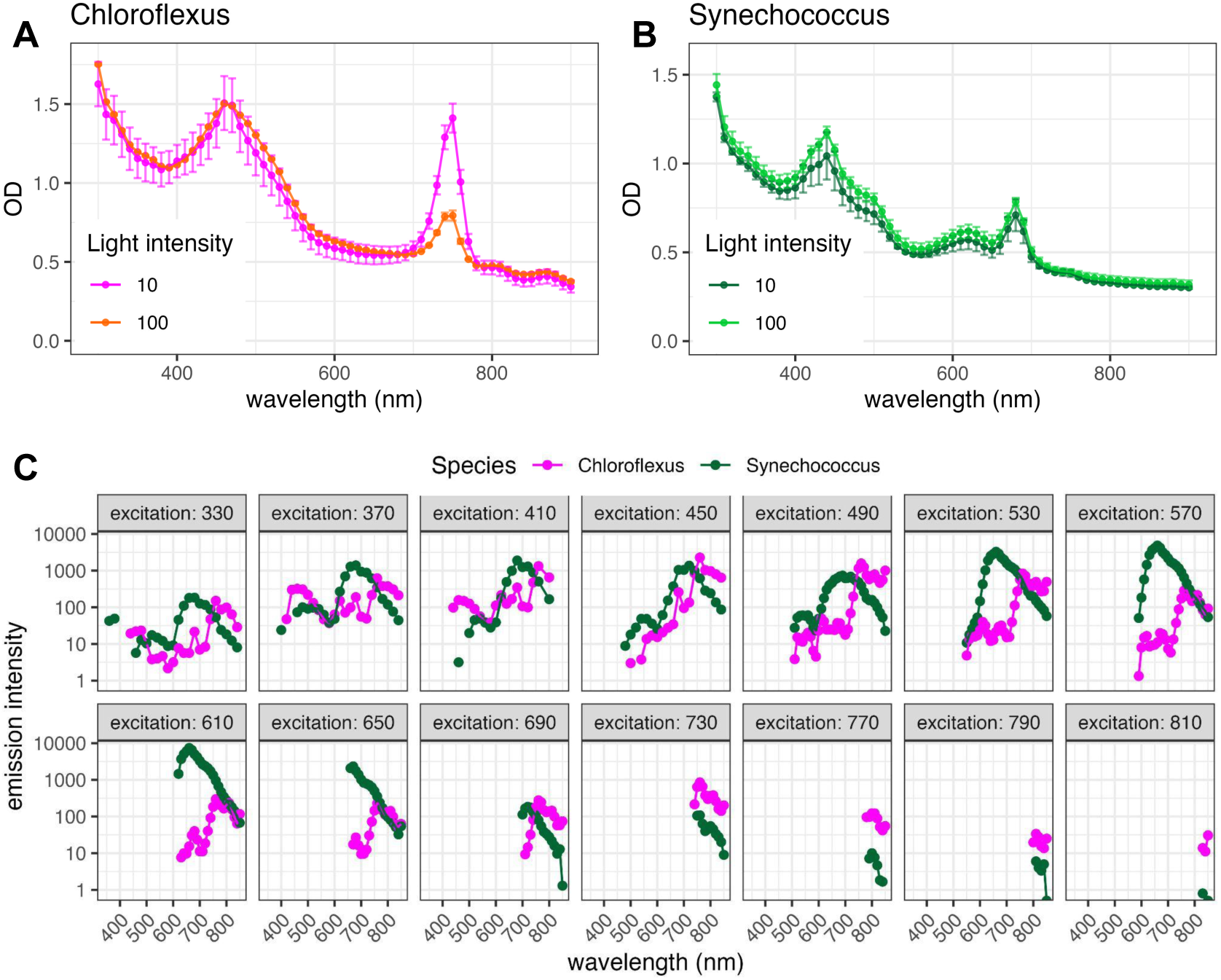
Absorption and fluorescence emission spectra of Chloroflexus (Chfl MS-1) and Synechococcus (Syn OS-B’) cultures. **(A)&(B)** Absorption spectra of Syn OS-B’ and Chfl MS-1 grown at 10 or 100 μmol·m^−2^·s^−1^ in dark and light colours respectively. **(C)** Fluorescence emission spectra of Chfl MS-1 (orange) and Syn OS-B’ (green) after excitation at different wavelengths indicated above each panel. Points represent means, error bars represent SD, n=3.

**Figure S3.**
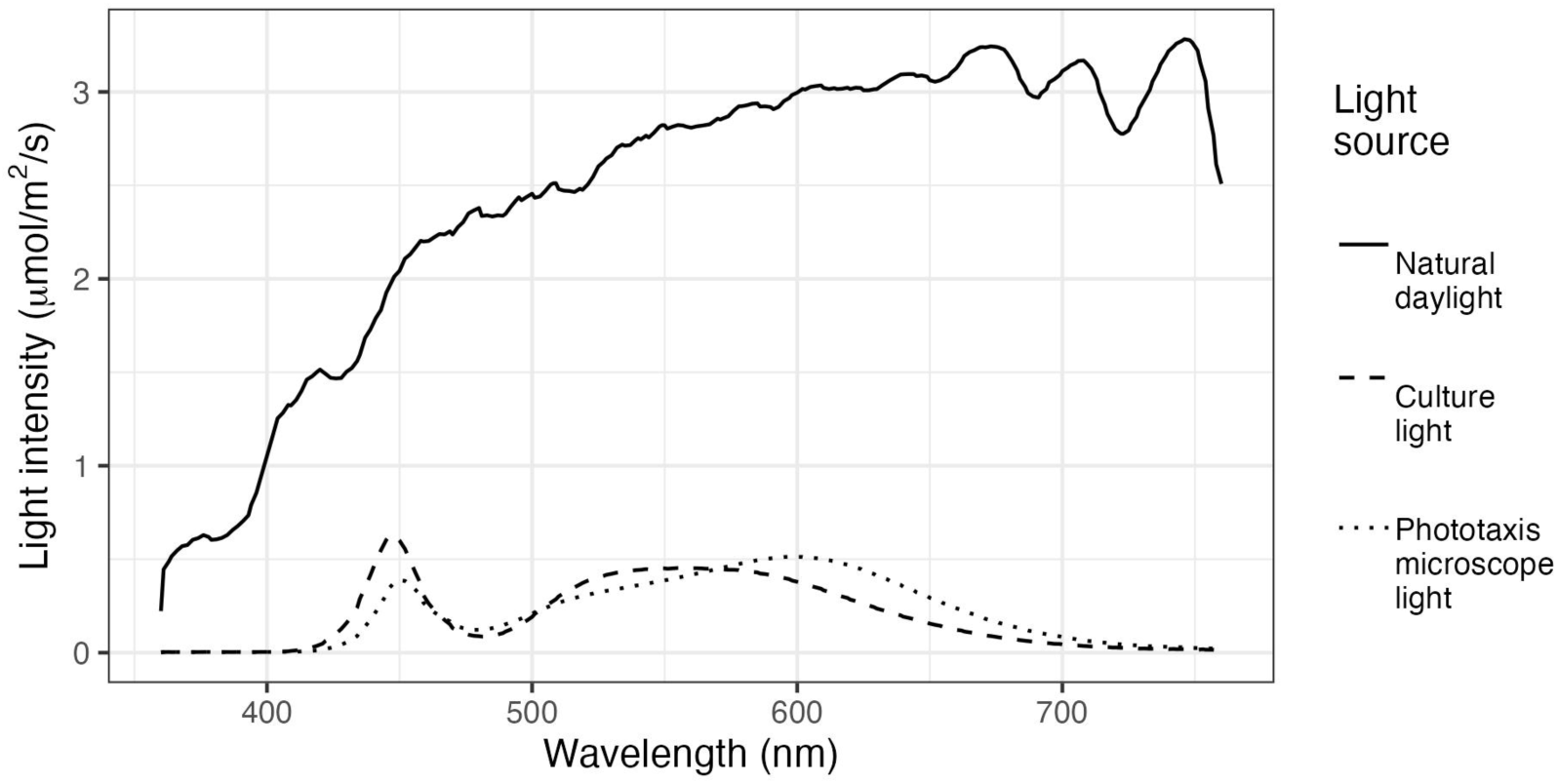
Light spectra of different light sources. Light spectra were measured with a Hipoint spectrophotometer between 360 and 760 nm. Natural daylight was recorded vertically at 3pm on the 03/25/21 (a partly sunny day) at 37°25’42.8"N 122°10’47.0"W. Culture light: spectrum during preculturing and collective motility assays (measured at the nearest edge of the culture vessels). Phototaxis microscope light: spectrum when measuring single cell phototactic/motility responses in a temperature-controlled microscope at the same point as the cells.

**Figure S4.**
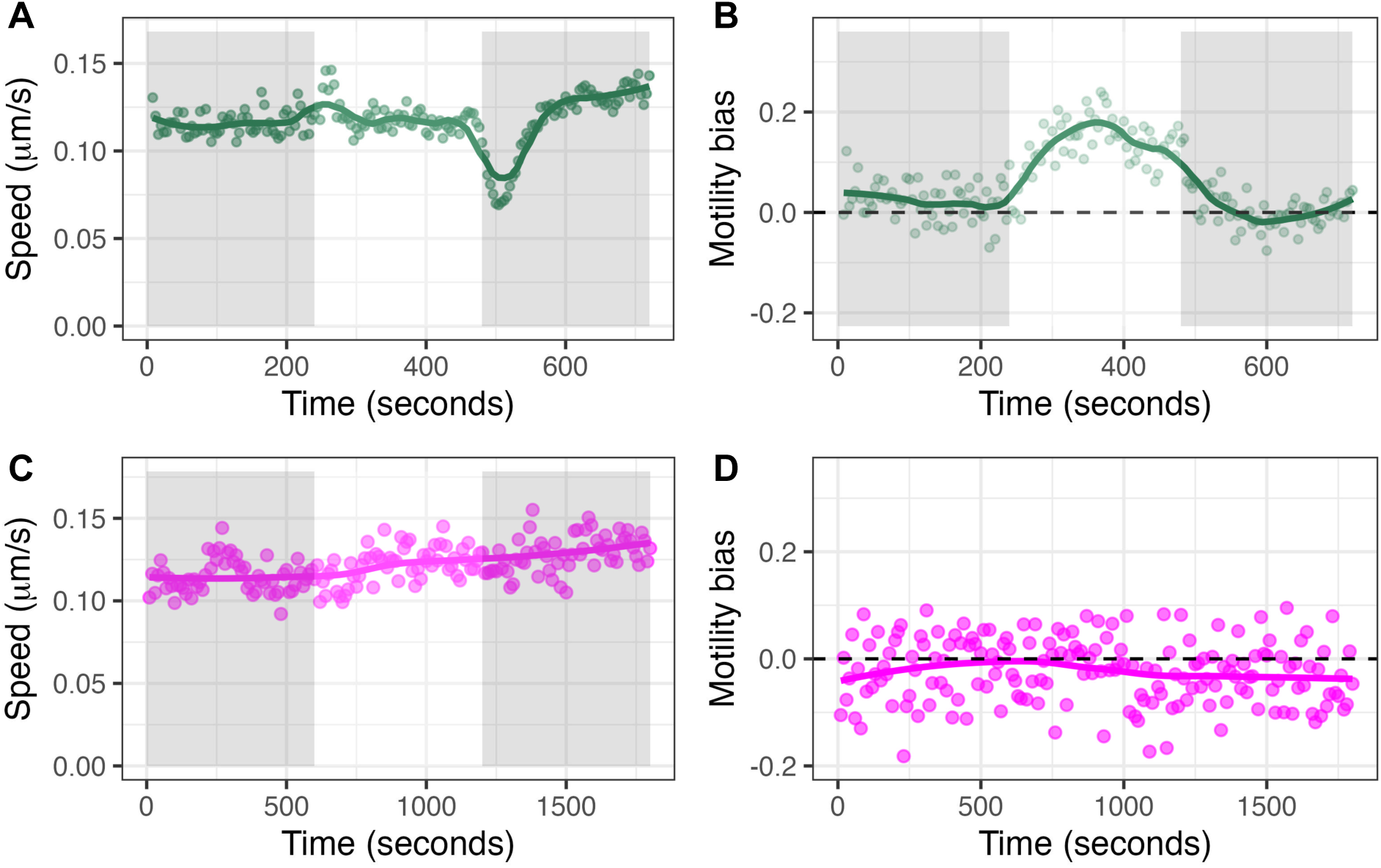
Syn OS-B’ cells and Chfl MS-1 filaments move differently during light transitions. Syn OS-B’ and Chfl MS-1 were both plated on DH10 solidified with 0.4% agarose, but Syn OS-B’ was incubated under directional white light for 2-4 days prior to tracking cells within phototactic projections, while Chfl MS-1 was incubated for 2-6 hours and filament tips tracked from within the starting droplet area. Cells were incubated in the microscope in the dark for >5 minutes then recorded under 4 or 10 minute periods (Syn OS-B’ and Chfl MS-1, respectively) of darkness, then 50 µmol·m^-2^·s^-1^ white light from right, then darkness. Chfl MS-1 filaments were tracked with a custom tip-tracking algorithm (see methods) and Syn OS-B’ cells by ImageJ using the Weka trainable segmentation and Trackmate plugins. Note that data in panels A&B are also shown in Figure 3 H&I. **(A)** Speed of tracked Syn OS-B’ cells within phototactic projections during light transitions. **(B)** Motility bias (proportion of movement towards light. 0 = random) of tracked Syn OS-B’ cells within phototactic projections during light transitions. **(C)** Speed of tracked Chfl MS-1 filaments. **(D)** Motility bias of Chfl MS-1 filaments. Each point represents the mean of >100 Chfl MS-1 tips or >300 Syn OS-B’ cells from 8-10 separate videos.

**Figure S5.**
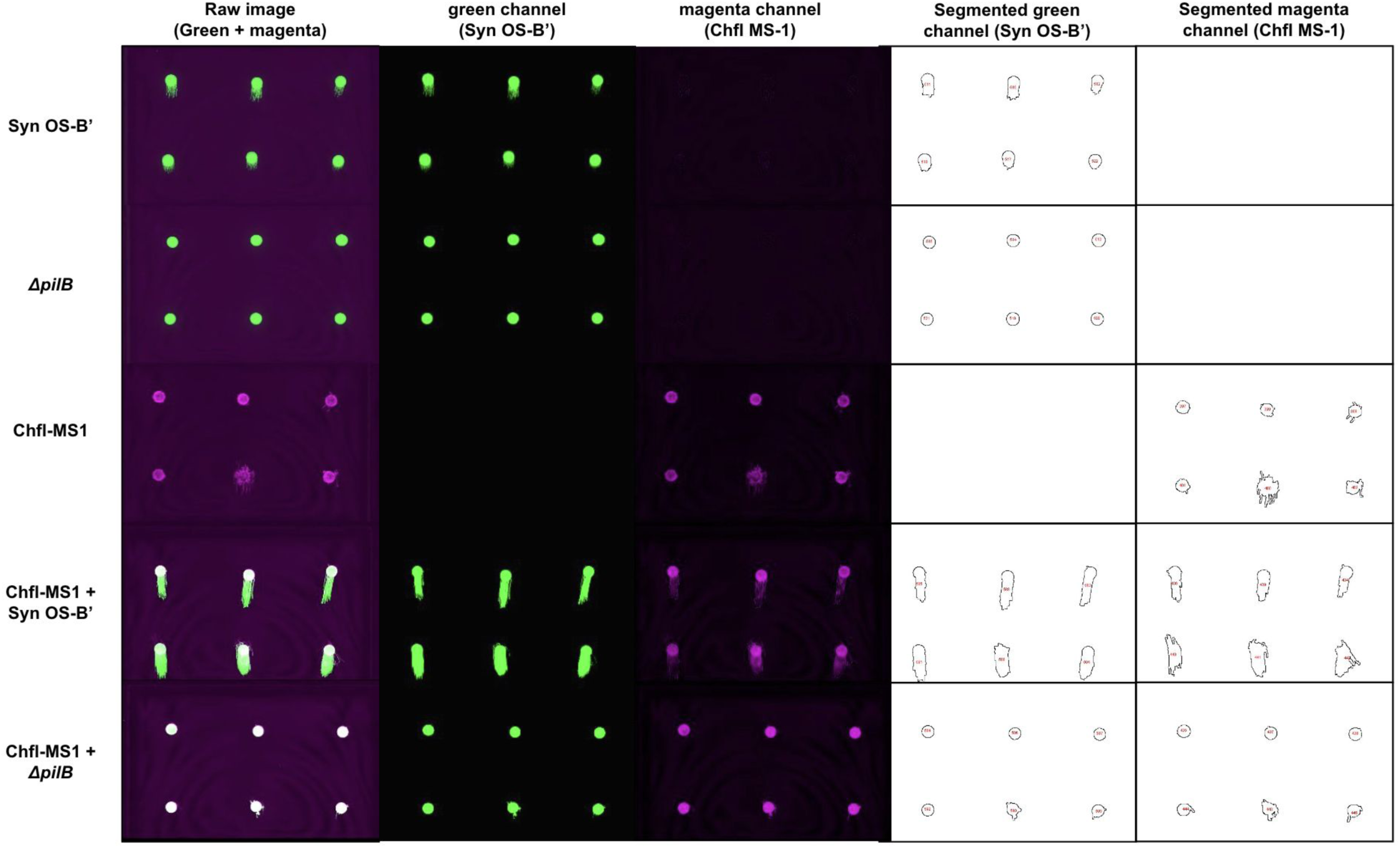
Visual overview of quantifying collective movement of consortia. Rectangular motility plates with 6 biological replicates (strains cultured separately for 3 months) were incubated for 4 days under directional light then imaged. Excitation was 784 nm and emission at 832BP37 nm (magenta) and excitation at 658 nm and emission at 710BP40 nm (green). A Fiji script was written to separate green and magenta channels and then threshold the images to select areas above the threshold. The pipeline was created using preliminary experiments and then applied without modification to our test data. The thresholded and selected images are shown in columns 2 and 3. Areas and center of mass coordinates were calculated for the objects and matched to the metadata for the experiment layout using the Hungarian algorithm. This allowed us to then plot the change in area or distance moved towards the light (in main text) by comparing processed scans at different timepoints.

**Figure S6.**
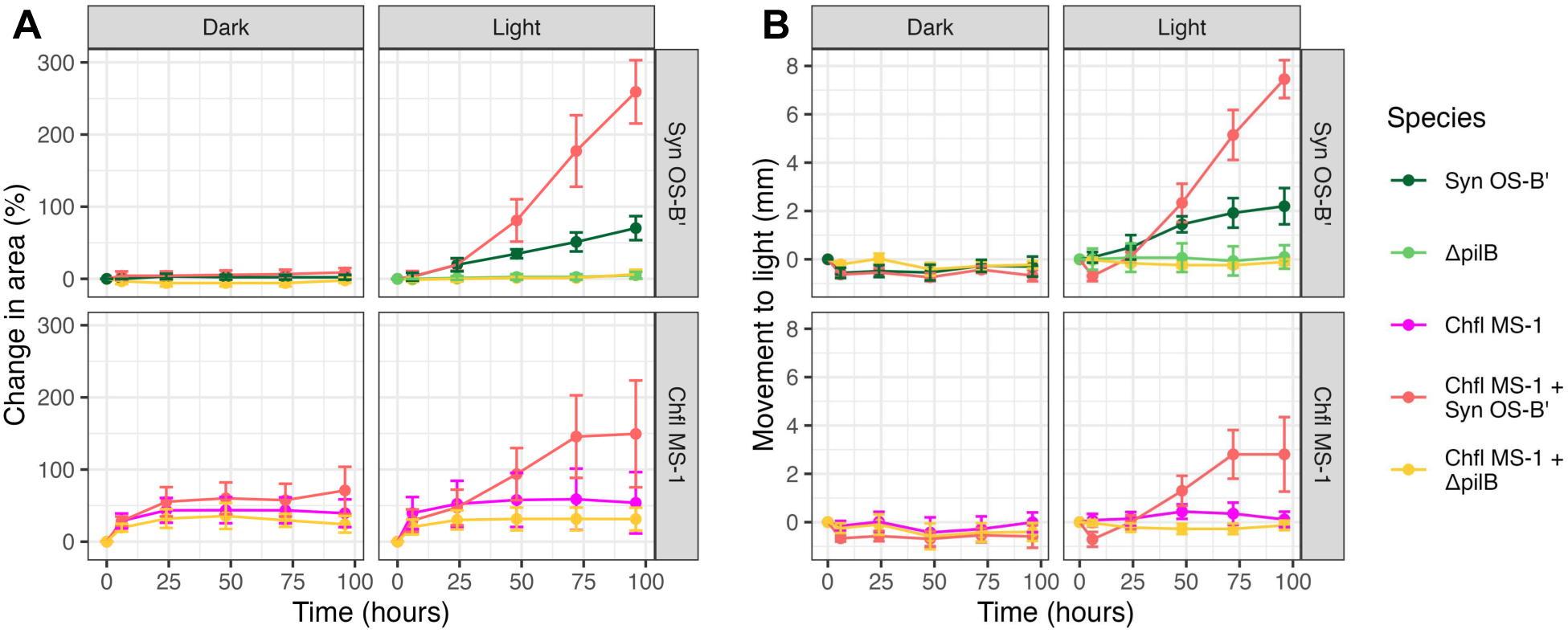
Syn OS-B’-Chfl MS-1 interactions improve movement towards the light. **(A)** Change in area of colonies. **(B)** Movement towards the light source of species mixtures. Syn OS-B’ chlorophyll signal (top panels) and Chfl MS-1 bacteriochlorophyll signal (bottom panels) in the dark (left panels) and directional 50 µmol·m^-2^·s^-1^ white light (right panel), with different species mixtures indicated by colours listed in the legend to the right. N=12 biological replicates from 2 experiments, and error bars indicate 95% confidence interval of the mean.

**Figure S7.**
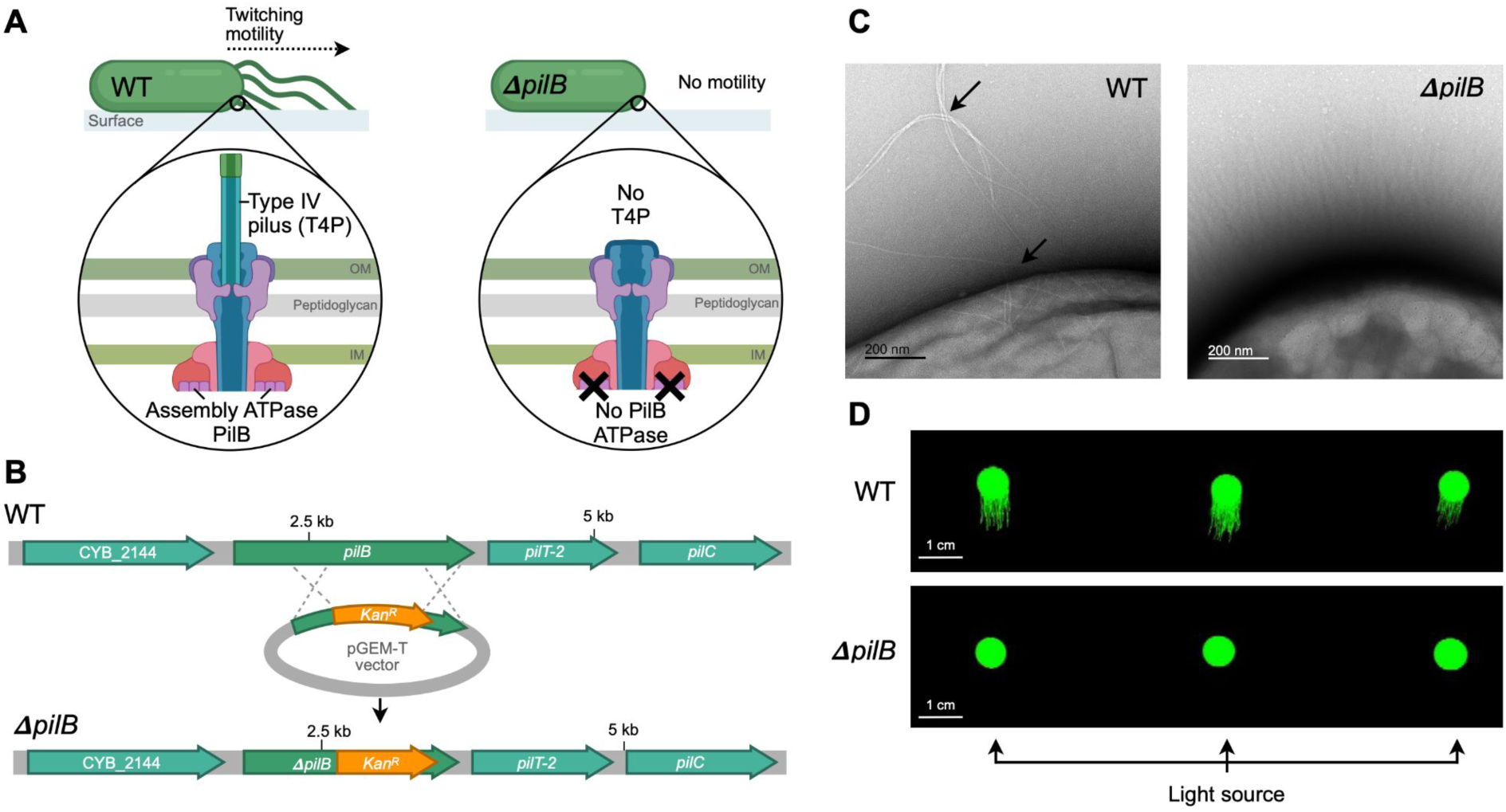
Syn OS-B’ *ΔpilB* mutants lack Type IV pili and motility. **(A)** Schematic of the intended effect of knocking out pilB in Syn OS-B’: the cells become non motile due to the lack of T4P, which is due to an inability to assemble pilin subunits into a pilus. OM, outer membrane; IM, inner membrane. **(B)** Diagram of the protocol by which a plasmid containing a Kanamycin resistance cassette (KanR) flanked by *pilB* sequences allowed integration into the *pilB* gene by homologous recombination using natural transformation of Syn OS-B’. **(C)** Transmission electron microscopy of Syn OS-B’ (WT (left) and *ΔpilB* (right)) shows T4P pili only in WT (indicated by black arrows), and regions of either EPS or thin pili in both (indicated by white arrows). Additional images are provided in Figures S5 and S6. **(D)** Chlorophyll fluorescence scans of 3 representative replicates of Syn OS-B’ cells (top) and independent *ΔpilB* knockouts (bottom) incubated under directional (from bottom) white light for 4 days, showing phototaxis of Syn OS-B’ but not *ΔpilB*, towards the light. Image in A created with BioRender.com.

**Figure S8.**
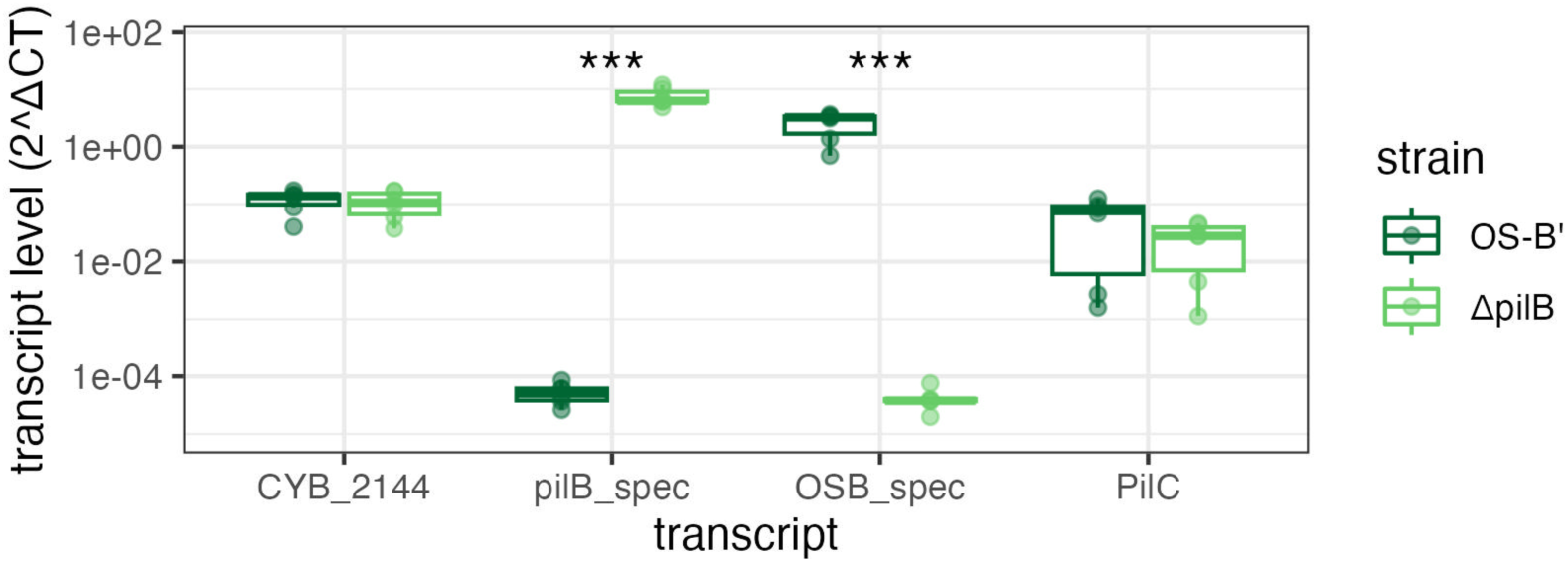
RT-qPCR on Syn OS-B’ and *ΔpilB* mutants reveals complete knockout of *pilB* gene and maintained expression of other genes in the operon. RNA was extracted from six late log-phase cultures of Syn OS-B’ and from six independent *ΔpilB* mutant cultures, reverse transcribed, and cDNA was quantified by qPCR. Primers were designed to amplify <300bp regions from the gene 5’ to *pilB* (CYB_2144), the kanamycin resistance cassette found only in *ΔpilB* (pilB_spec), the region of *pilB* that was knocked out and therefore only present in Syn OS-B’ (OSB_spec), and *pilC*, which is 3’ to *pilB* and possibly in the same operon. Transcript concentration is displayed on the y-axis and calculated by raising 2 to the power of the the threshold cycle value of the target gene subtracted from the mean threshold cycle of four housekeeping genes (cysE, La, U32, SufS) (2^-ΔCT^ method). Student’s t-tests were performed with Bonferroni correction with statistical significance reported as follows: *** = p<0.001, otherwise p>0.05.

**Figure S9.**
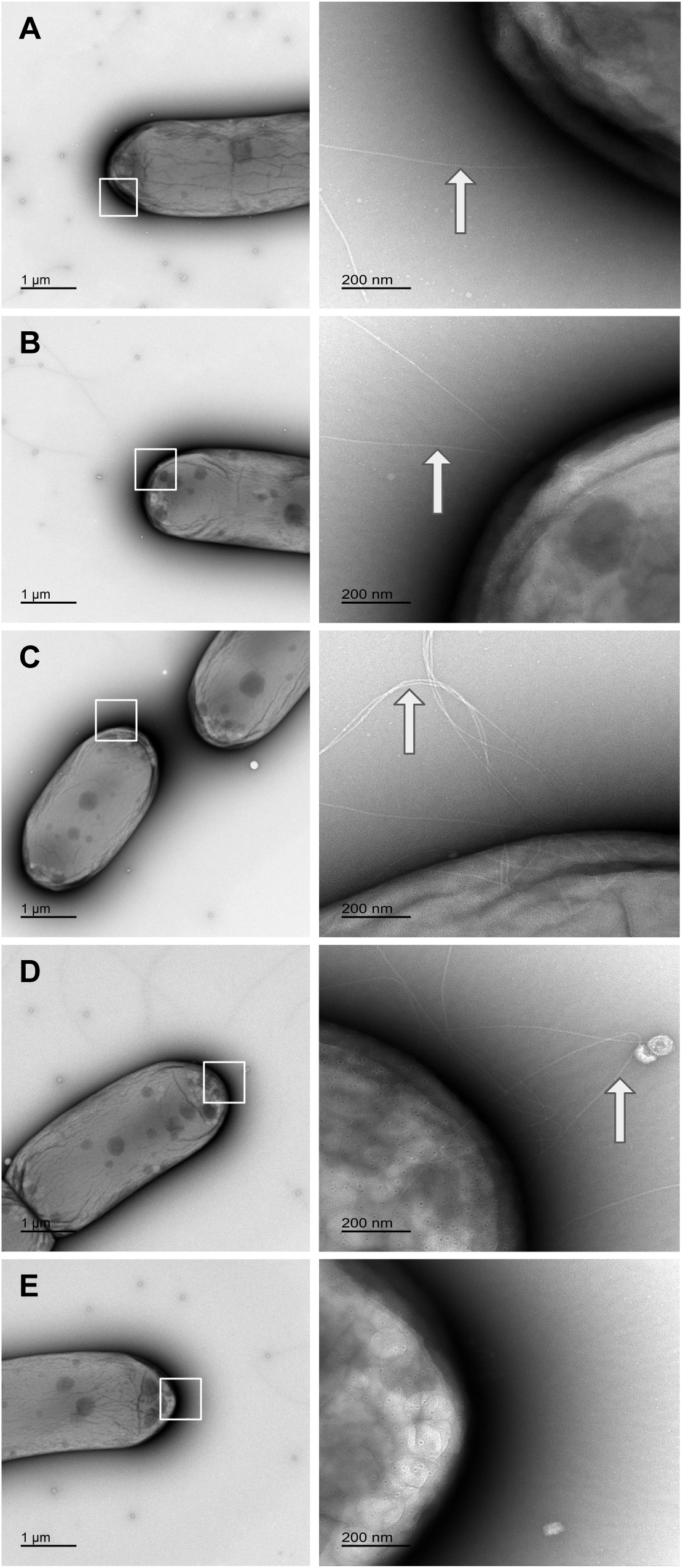
Transmission electron microscopy of Syn OS-B’ cells reveals thick pili (indicated by arrows). Cells were stained with uranyl acetate and mounted on TEM grids prior to imaging. 5 cells were randomly selected and a magnified image of one of their poles was captured. 4 out of 5 cells had thick pili near their poles.

**Figure S10.**
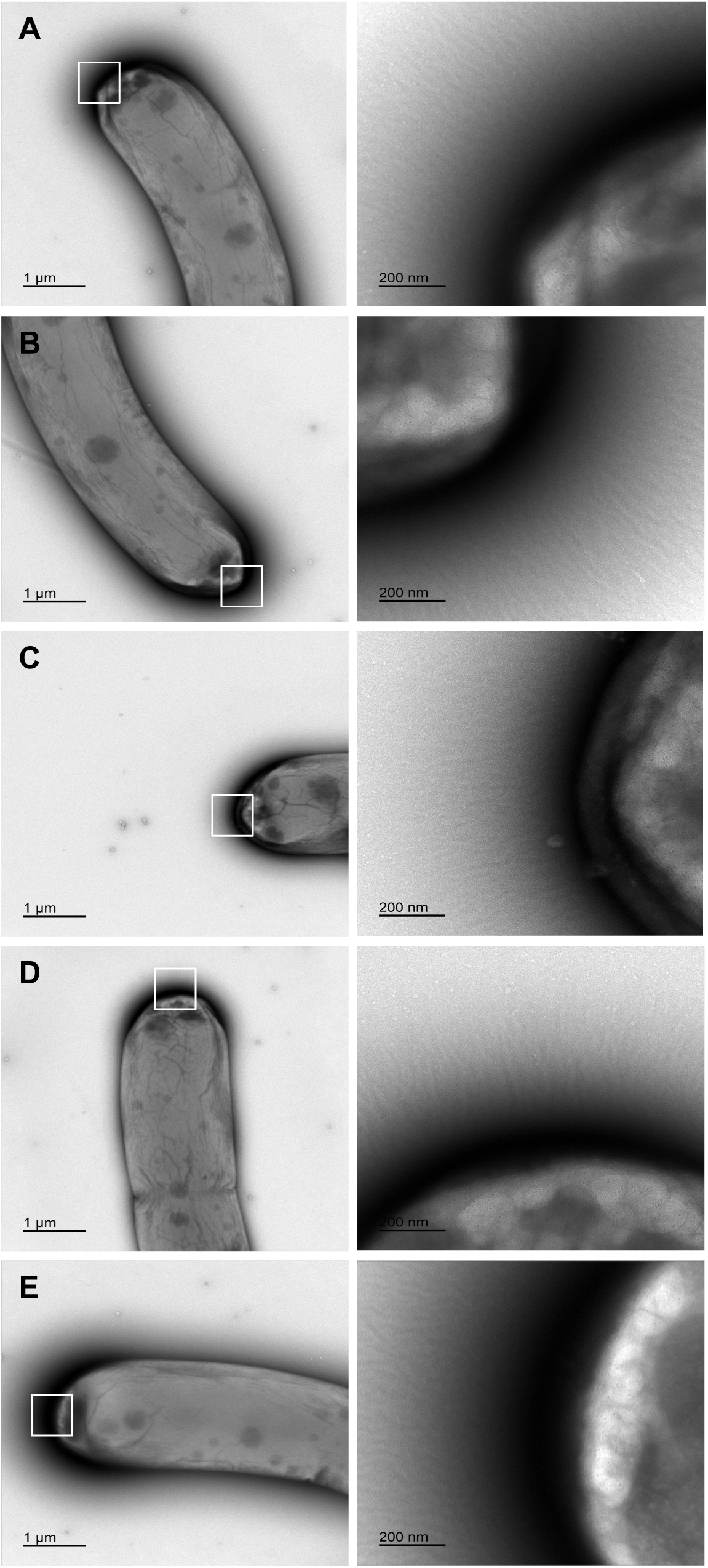
Transmission electron microscopy of *ΔpilB* cells reveals absence of thick pili. Cells were stained with uranyl acetate and mounted on TEM grids prior to imaging. Five cells were randomly selected and a magnified image of one of their poles was captured. 0 out of 5 cells had thick pili near their poles.

**Figure S11.**
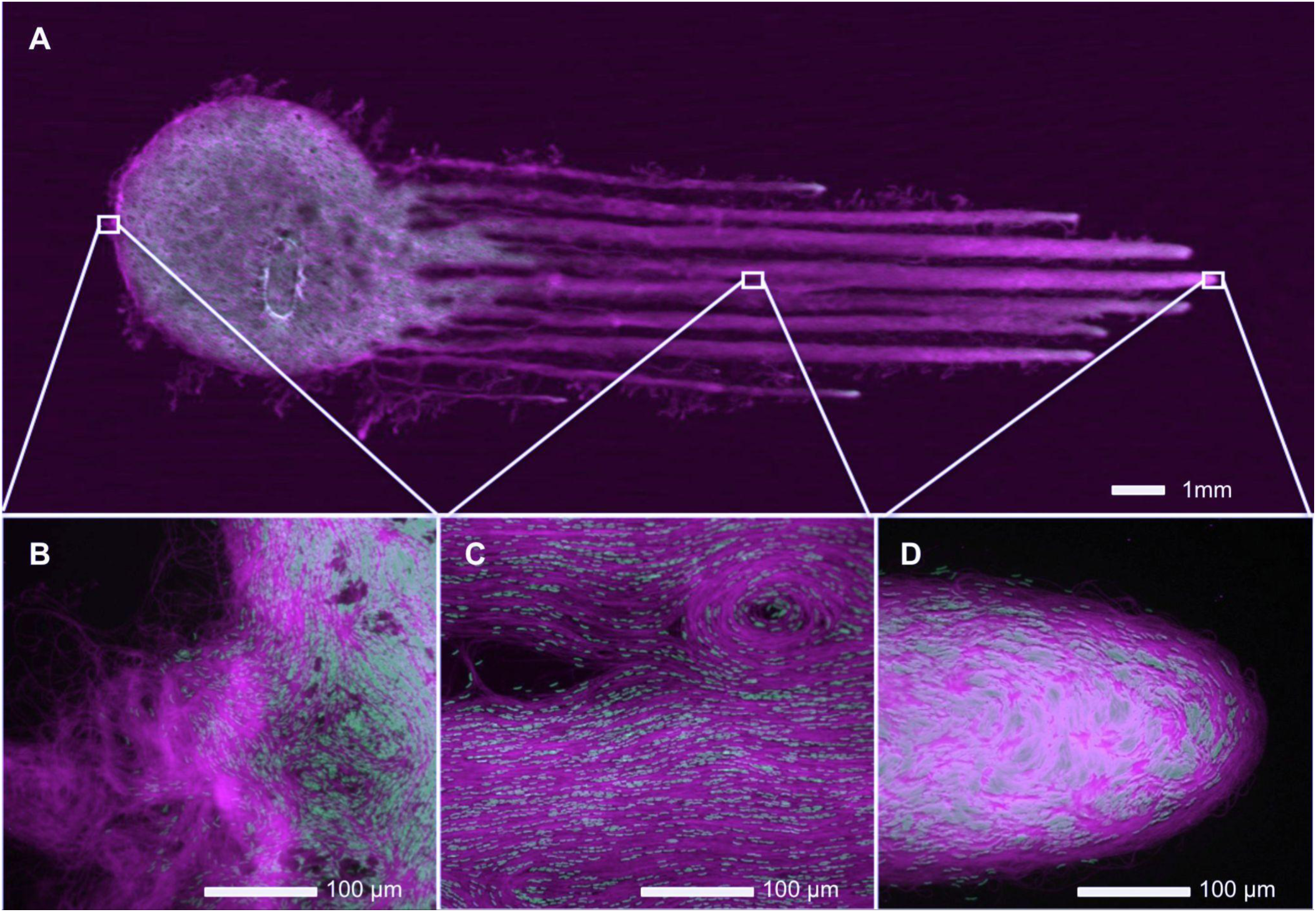
Viewing Chfl MS-1-Syn OS-B’ consortium motility with fluorescence scanning and microscopy. Images were taken after 4 days incubation under directional (from right) 60 µmol·m^-2^·s^-1^ white light. **(A)** Fluorescence microscopy with excitation 784 nm and emission at 832 nm (band pass of 37 nm) (magenta) and by excitation 658 nm and emission at 710 nm (band pass of 40 nm) (green). **(B,C,D)** Fluorescence micrographs of Chfl MS-1 + Syn OS-B’ from selected regions (squares) with excitation at 450-490 nm and emission at >515 nm (magenta), and excitation at 540-580 nm and emission at 608-682 nm (green).

**Figure S12.**
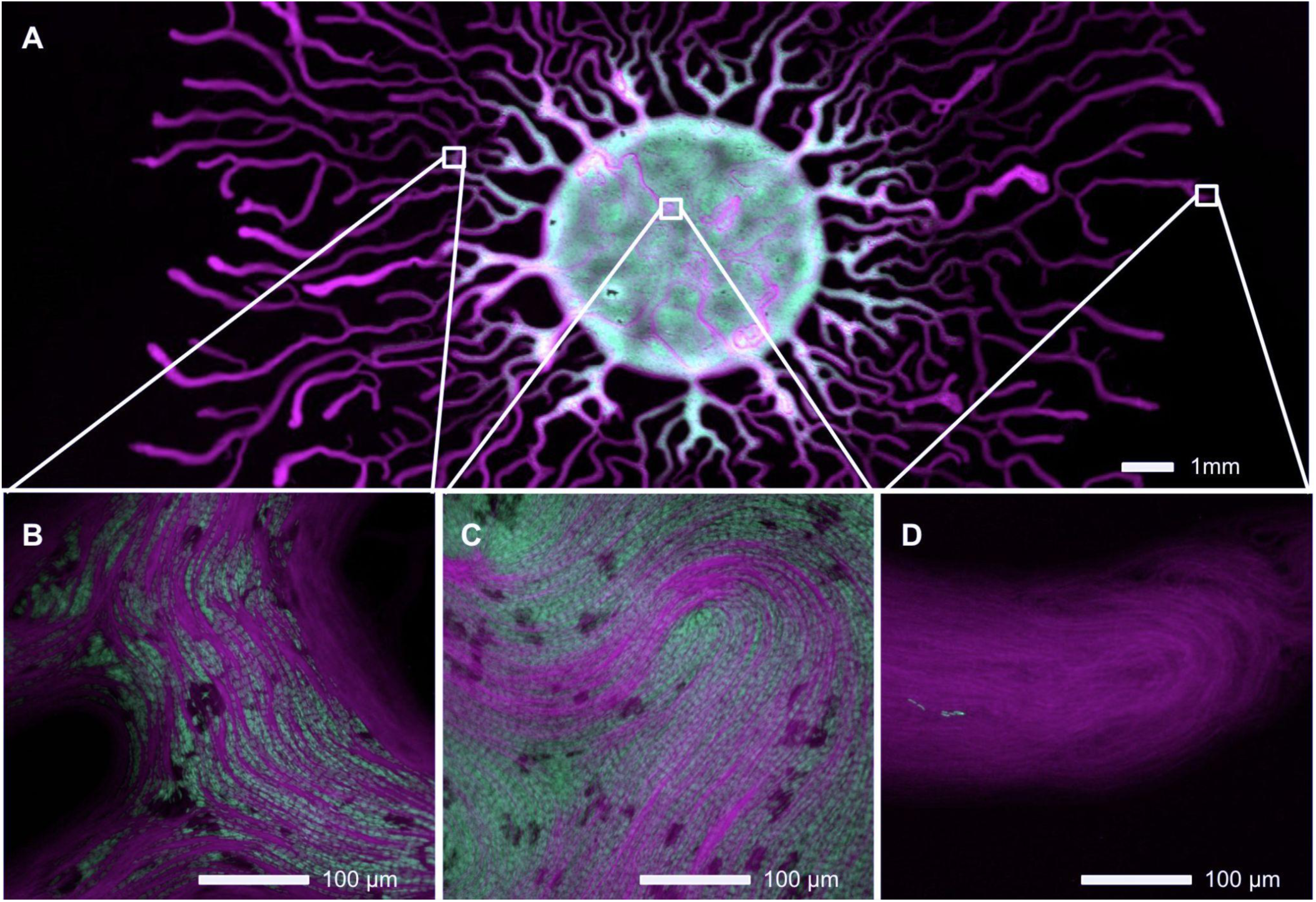
Viewing Chfl MS-1-*ΔpilB* consortium motility with fluorescence scanning and microscopy. Images were taken after 4 days incubation under directional (from right) 60 µmol·m^-2^·s^-1^ white light. **(A)** Fluorescence microscopy with excitation 784 nm and emission at 832 nm (band pass of 37 nm) (magenta) and by excitation 658 nm and emission at 710 nm (band pass of 40 nm) (green). **(B,C,D)** Fluorescence micrographs of Chfl MS-1 + *ΔpilB* from selected regions (squares) in panel A with excitation at 450-490 nm and emission at >515 nm (magenta), and excitation at 540-580 nm and emission at 608-682 nm (green).

**Figure S13.**
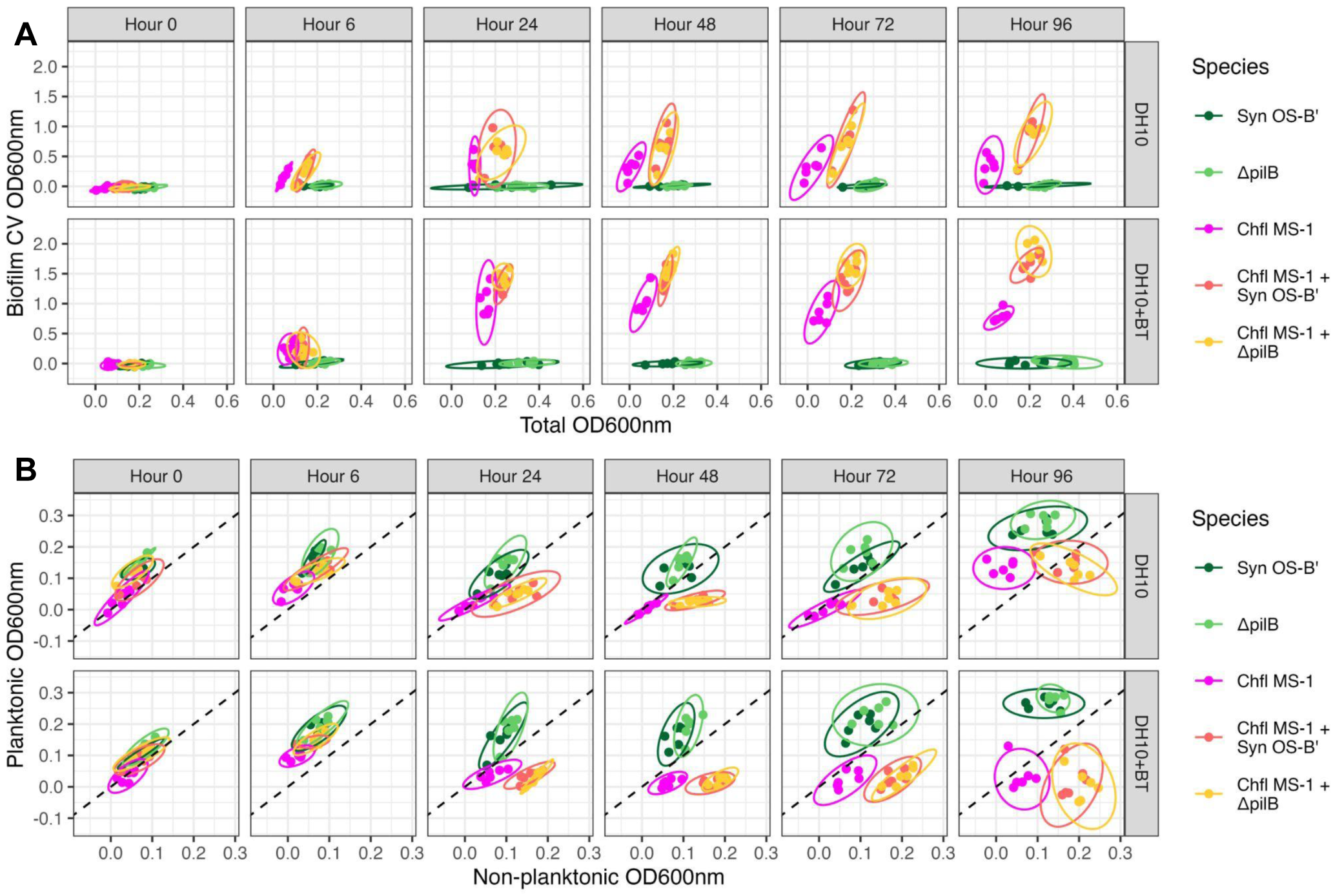
Binary consortia consistently develop more biofilm and less planktonic cells than axenic cyanobacteria over time. **(A)** After 0-4 days of incubation of cultures, OD600nm absorption measurements were taken on the total culture (x-axis) and after crystal violet staining of the biofilm (y-axis). **(B)** After 0-4 days, cultures were briefly disrupted by pipetting and 80% of the culture volume was transferred to a new plate. OD600nm was measured in the 20% of the remaining volume and called “Non-planktonic OD600nm” (x-axis), and in the transferred volume and called “Planktonic OD600nm” (y-axis). Top panel labels indicate the time of sampling after inoculation, side panel labels indicate culture medium (DH10 or DH10 + BT (bactotryptone)). Colours represent the species in cultures, as indicated by the legend to the right, and ellipses indicate the 95% confidence intervals of the measurements. N=6 independent biological replicates for every timepoint and medium.

**Figure S14.**
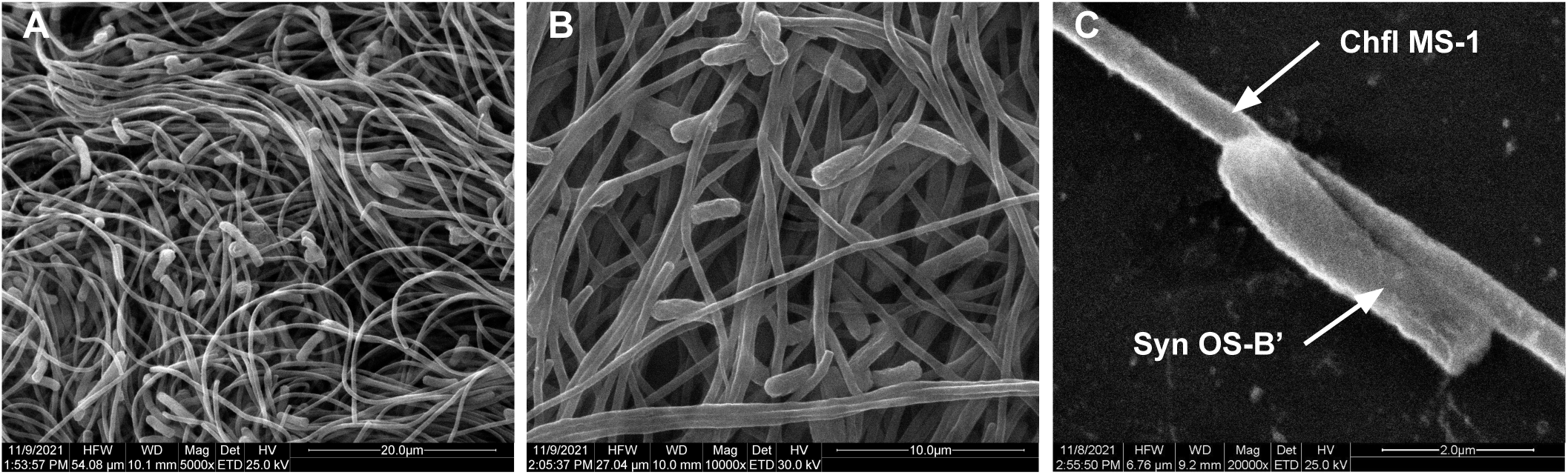
Scanning electron microscopy of Syn OS-B’ and Chfl MS-1 binary consortium reveals that the species are often associated with one another. The samples were imaged in Quanta 200 (FEI) SEM with 25 kV accelerating voltage in high vacuum with secondary emission (SE) detector. Increasing magnification from left to right **(A)** 5000x, **(B)** 10,000x, **(C)** 20,000x.

**Figure S15.**
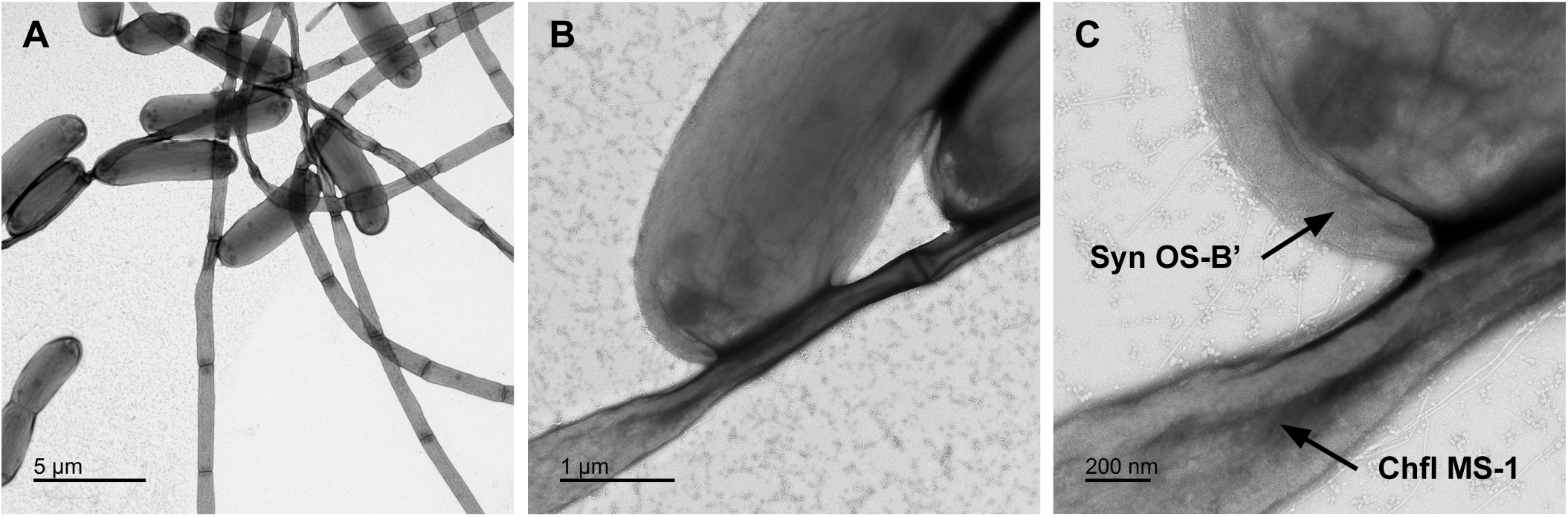
Transmission electron microscopy of Syn OS-B’ and Chfl MS-1 binary consortium reveals that the species are often associated with one another. The grids with uranyl-acetate stained cells were imaged in JEOL JEM-1400 with 120 kV accelerating voltage equipped with a Gatan Inc. OneView 4kX4k sCMOS camera. Increasing magnification from left to right **(A)** 5,000x, **(B)** 30,000x, **(C)** 80,000x.

**Table S1.**
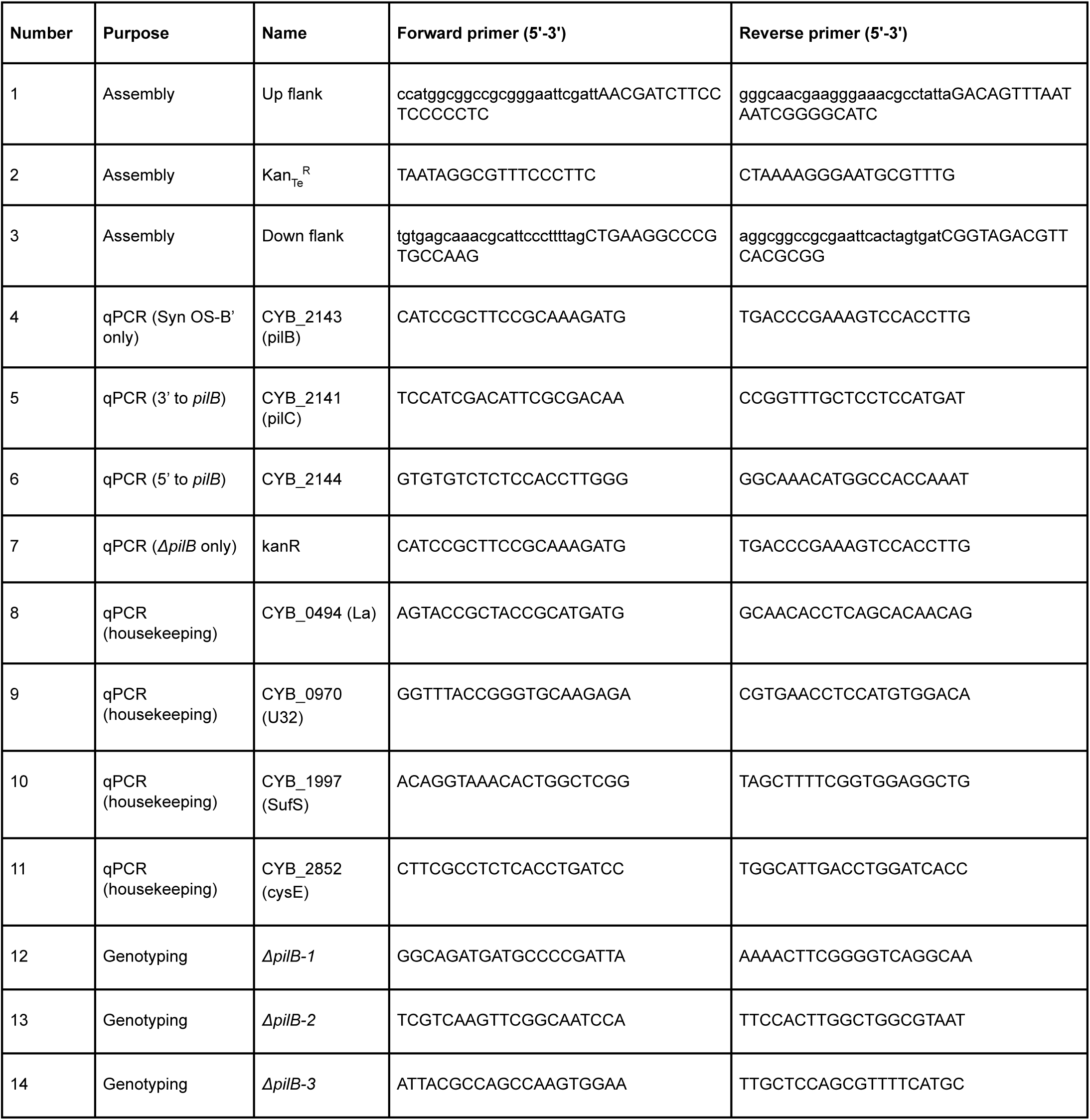
Primers used in this study. Primers were used to create the construct used for transformation (Assembly), test whether *pilB* was replaced by the cassette in *ΔpilB* (Genotyping), and to test for a polar effect of the kanamycin resistance cassette insertion by quantifying relative transcript levels of nearby genes (qPCR).

## Supplementary Video legends

**Video S1. Collective movement of Syn OS-B’ towards white light (from the right).** Video duration: 10 hours. Video speed: 6000x (100 minutes per second). Magnification: 40x. Species: Synechococcus sp. JA-2-3B’a(2-13). Surface: DH10 medium solidified with 0.4% agarose. Temperature: 50℃. Light: 20 µmol·m^-2^·s^-1^ white light from right. Time in hours and minutes shown in top left.

**Video S2. Collective movement of Chfl MS-1 in darkness.** Video duration: 2 hours. Video speed: 1200x (20 minutes per second). Magnification: 40x. Species: Chloroflexus MS-CIW1. Surface: DH10 medium solidified with 0.4% agarose. Temperature: 50℃. Light: Darkness. Time in hours and minutes shown in top left.

**Video S3. Movement of single Syn OS-B’ cells under dynamic light conditions.** Syn OS-B’ cells are viewed in the middle of a phototactic projection over a 12 minute period starting in the dark, then with white light at 50 µmol·m^-2^·s^-1^ from right hand side switched on at 4 minutes and off at 8 minutes. Syn OS-B’ cells were identified and outlined using Trackmate in ImageJ. Video speed: 72x (1.2 minutes per second). Magnification: 200x. Species:

Synechococcus OS-B’. Surface: DH10 medium solidified with 0.4% agarose. Temperature: 50℃. Light condition and direction with time in seconds shown in top left.

**Video S4. Movement of single Chfl MS-1 filaments under dynamic light conditions.** Chfl MS-1 filaments were viewed 2 hours after plating on DH10 agarose media over a 30 minute period starting in the dark, then with white light at 50 µmol·m^-2^·s^-1^ from right hand side switched on at 10 minutes and off at 20 minutes. Chfl MS-1 filaments were identified and outlined using ridge detection in ImageJ. Video speed: 180x (3 minutes per second).

Magnification: 200x. Species: Chloroflexus MS-CIW-1. Surface: DH10 medium solidified with 0.4% agarose. Temperature: 50℃. Light condition and direction with time in seconds shown in top left.

**Video S5. Collective movement of Chfl MS-1 (left) and Syn OS-B’ (right) axenically and when mixed together (middle).** Cultures were illuminated with 50 µmol·m^-2^·s^-1^ white light from the bottom of the image and scanned at intervals of several hours with excitation 784 nm and emission at 832BP37 nm (magenta) and by excitation 658 nm and emission at 710BP40 nm (green). Time in hours is displayed in top left.

**Video S6. Collective movement of Syn OS-B’ and Chfl MS-1 towards white light (from the right).** Video duration: 4 hours. First hour in the dark, then white light from right is switched on at 1 hour. Video speed: 1200x (20 minutes per second). Magnification: 40x. Surface: DH10 medium solidified with 0.4% agarose. Temperature: 50℃. Light: 50 µmol·m^-2^·s^-1^ white light. Light condition and direction with time in minutes shown in top left.

**Video S7. Collective movement of ΔpilB and Chfl MS-1 with white light (from the right).** Video duration: 4 hours. First hour in the dark, then white light from right is switched on at 1 hour. Video speed: 1200x (20 minutes per second). Magnification: 40x. Surface: DH10 medium solidified with 0.4% agarose. Temperature: 50℃. Light: 50 µmol·m^-2^·s^-1^ white light. Light condition and direction with time in minutes shown in top left.

**Video S8. Movement of Syn OS-B’ cells within Chfl MS-1 bundles under dynamic light conditions.** Syn OS-B’ cells are viewed in the middle of a phototactic projection composed of Syn OS-B’ and Chfl MS-1 over a 12-minute period starting in the dark, then with white light from right hand side switched on at 4 minutes, and switched off at 8 minutes. Syn OS-B’ cells were identified and outlined using Trackmate in ImageJ. Video speed: 72x (1.2 minutes per second). Magnification: 200x. Species: Syn OS-B’. Surface: DH10 medium solidified with 0.4% agarose.

Temperature: 50℃. Light condition and direction with time in seconds shown in top left.

**Video S9. Movement of *ΔpilB* cells within Chfl MS-1 bundles under dynamic light conditions.** ΔpilB cells are viewed in the middle of a projection composed of ΔpilB and Chfl MS-1 over a 12-minute period starting in the dark, then with white light from right hand side switched on at 4 minutes, and switched off at 8 minutes. ΔpilB cells were identified and outlined using Trackmate in ImageJ. Video speed: 72x (1.2 minutes per second). Magnification: 200x. Species: ΔpilB Syn OS-B’. Surface: DH10 medium solidified with 0.4% agarose. Temperature: 50℃. Light condition and direction with time in seconds shown in top left.

## References

[1] W.-S. Shu and L.-N. Huang, “Microbial diversity in extreme environments,” Nat. Rev. Microbiol., pp. 1–17, Nov. 2021.

[2] L. J. Stal, “Physiological ecology of cyanobacteria in microbial mats and other communities,” New Phytol., vol. 131, no. 1, pp. 1–32, Sep. 1995.

[3] S. I. Jensen, A.-S. Steunou, D. Bhaya, M. Kühl, and A. R. Grossman, “In situ dynamics of O2, pH and cyanobacterial transcripts associated with CCM, photosynthesis and detoxification of ROS,” ISME J., vol. 5, no. 2, pp. 317–328, Feb. 2011.

[4] J. M. Keegstra, F. Carrara, and R. Stocker, “The ecological roles of bacterial chemotaxis,” Nat. Rev. Microbiol., vol. 20, no. 8, pp. 491–504, Aug. 2022.

[5] A. Wilde and C. W. Mullineaux, “Light-controlled motility in prokaryotes and the problem of directional light perception,” FEMS Microbiol. Rev., vol. 41, no. 6, pp. 900–922, Nov. 2017.

[6] K. Sauer et al., “The biofilm life cycle: expanding the conceptual model of biofilm formation,” Nat. Rev. Microbiol., vol. 20, no. 10, pp. 608–620, Oct. 2022.

[7] P. Chiang and L. L. Burrows, “Biofilm formation by hyperpiliated mutants of Pseudomonas aeruginosa,” J. Bacteriol., vol. 185, no. 7, pp. 2374–2378, Apr. 2003.

[8] M. Klausen, A. Aaes-Jørgensen, S. Molin, and T. Tolker-Nielsen, “Involvement of bacterial migration in the development of complex multicellular structures in Pseudomonas aeruginosa biofilms,” Mol. Microbiol., vol. 50, no. 1, pp. 61–68, Oct. 2003.

[9] C. Li et al., “Social motility of biofilm-like microcolonies in a gliding bacterium,” Nat. Commun., vol. 12, no. 1, p. 5700, Sep. 2021.

[10] K. Toida et al., “The GGDEF protein Dgc2 suppresses both motility and biofilm formation in the filamentous cyanobacterium Leptolyngbya boryana,” Microbiol Spectr, vol. 11, no. 5, p. e0483722, Sep. 2023.

[11] K. A. Brileya, L. B. Camilleri, G. M. Zane, J. D. Wall, and M. W. Fields, “Biofilm growth mode promotes maximum carrying capacity and community stability during product inhibition syntrophy,” Front. Microbiol., vol. 5, p. 693, Dec. 2014.

[12] L. Ma, M. E. Konkel, and X. Lu, “Antimicrobial Resistance Gene Transfer from Campylobacter jejuni in Mono- and Dual-Species Biofilms,” Appl. Environ. Microbiol., vol. 87, no. 15, p. e0065921, Jul. 2021.

[13] N. K. Ratheesh, A. M. Zdimal, C. A. Calderon, and A. Shrivastava, “Bacterial Swarm- Mediated Phage Transportation Disrupts a Biofilm Inherently Protected from Phage Penetration,” Microbiol Spectr, vol. 11, no. 4, p. e0093723, Aug. 2023.

[14] R. G. Perkins et al., “Vertical cell movement is a primary response of intertidal benthic biofilms to increasing light dose,” Mar. Ecol. Prog. Ser., vol. 416, pp. 93–103, Oct. 2010.

[15] J. M. Wood et al., “Assessing microbial diversity in Yellowstone National Park hot springs using a field deployable automated nucleic acid extraction system,” Frontiers in Ecology and Evolution, vol. 12, 2024, doi: 10.3389/fevo.2024.1306008.

[16] L. Steinke et al., “Short-Term Stable Isotope Probing of Proteins Reveals Taxa Incorporating Inorganic Carbon in a Hot Spring Microbial Mat,” Appl. Environ. Microbiol., vol. 86, no. 7, Mar. 2020, doi: 10.1128/AEM.01829-19.

[17] W. N. Doemel and T. D. Brock, “Structure, growth, and decomposition of laminated algal- bacterial mats in alkaline hot springs,” Appl. Environ. Microbiol., vol. 34, no. 4, pp. 433–452, Oct. 1977.

[18] M. Lichtenberg, P. Cartaxana, and M. Kühl, “Vertical migration optimizes photosynthetic efficiency of motile cyanobacteria in a coastal microbial mat,” Frontiers in Marine Science, 2020, [Online]. Available: https://www.frontiersin.org/articles/10.3389/fmars.2020.00359/full

[19] N. B. Ramsing, M. J. Ferris, and D. M. Ward, “Highly ordered vertical structure of Synechococcus populations within the one-millimeter-thick photic zone of a hot spring cyanobacterial mat,” Appl. Environ. Microbiol., vol. 66, no. 3, pp. 1038–1049, Mar. 2000.

[20] S. Yoshihara and M. Ikeuchi, “Phototactic motility in the unicellular cyanobacterium Synechocystis sp. PCC 6803,” Photochem. Photobiol. Sci., vol. 3, no. 6, pp. 512–518, Jun. 2004.

[21] P. Italia, V. K. Pallipuram, and D. D. Risser, “Dynamic localization of HmpF regulates type IV pilus activity and directional motility in the filamentous cyanobacterium Nostoc punctiforme,” Molecular, 2017, [Online]. Available: https://onlinelibrary.wiley.com/doi/abs/10.1111/mmi.13761

[22] S.-I. Fukushima, S. Morohoshi, S. Hanada, K. Matsuura, and S. Haruta, “Gliding motility driven by individual cell-surface movements in a multicellular filamentous bacterium Chloroflexus aggregans,” FEMS Microbiol. Lett., vol. 363, no. 8, Apr. 2016, doi: 10.1093/femsle/fnw056.

[23] V. A. Gaisin, R. Kooger, D. S. Grouzdev, V. M. Gorlenko, and M. Pilhofer, “Cryo-Electron Tomography Reveals the Complex Ultrastructural Organization of Multicellular Filamentous Chloroflexota (Chloroflexi) Bacteria,” Front. Microbiol., vol. 11, p. 1373, Jun. 2020.

[24] A. Fourçans et al., “Vertical migration of phototrophic bacterial populations in a hypersaline microbial mat from Salins-de-Giraud (Camargue, France),” FEMS Microbiol. Ecol., vol. 57, no. 3, pp. 367–377, Sep. 2006.

[25] E. D. Becraft, F. M. Cohan, M. Kühl, S. I. Jensen, and D. M. Ward, “Fine-scale distribution patterns of Synechococcus ecological diversity in microbial mats of Mushroom Spring, Yellowstone National Park,” Appl. Environ. Microbiol., vol. 77, no. 21, pp. 7689–7697, Nov. 2011.

[26] L. Wörmer et al., “A micrometer-scale snapshot on phototroph spatial distributions: mass spectrometry imaging of microbial mats in Octopus Spring, Yellowstone National Park,” Geobiology, vol. 18, no. 6, pp. 742–759, Nov. 2020.

[27] N. B. Ramsing, M. J. Ferris, and D. M. Ward, “Light-induced motility of thermophilic Synechococcus isolates from Octopus Spring, Yellowstone National Park,” Appl. Environ. Microbiol., vol. 63, no. 6, pp. 2347–2354, Jun. 1997.

[28] J. P. Allewalt, M. M. Bateson, N. P. Revsbech, K. Slack, and D. M. Ward, “Effect of temperature and light on growth of and photosynthesis by Synechococcus isolates typical of those predominating in the octopus spring microbial mat community of Yellowstone National Park,” Appl. Environ. Microbiol., vol. 72, no. 1, pp. 544–550, Jan. 2006.

[29] J. Komárek, J. R. Johansen, J. Šmarda, and O. Strunecký, “Phylogeny and taxonomy of Synechococcus-like cyanobacteria,” Fottea, vol. 20, no. 2, pp. 171–191, Oct. 2020.

[30] A. N. Shelton et al., “Draft genome of Chloroflexus sp. MS-CIW-1, of the Chloroflexus sp. MS-G group from Mushroom Spring, Yellowstone National Park,” Microbiol Resour Announc, vol. 13, no. 3, p. e0071023, Mar. 2024.

[31] A. C. Bennett, S. K. Murugapiran, and T. L. Hamilton, “Temperature impacts community structure and function of phototrophic Chloroflexi and Cyanobacteria in two alkaline hot springs in Yellowstone National Park,” Environ. Microbiol. Rep., vol. 12, no. 5, pp. 503–513, Oct. 2020.

[32] F. Bunbury, C. Rivas, V. Calatrava, A. N. Shelton, A. Grossman, and D. Bhaya, “Differential Phototactic Behavior of Closely Related Cyanobacterial Isolates from Yellowstone Hot Spring Biofilms,” Appl. Environ. Microbiol., vol. 88, no. 10, p. e0019622, May 2022.

[33] M. Burriesci and D. Bhaya, “Tracking phototactic responses and modeling motility of Synechocystis sp. strain PCC6803,” J. Photochem. Photobiol. B, vol. 91, no. 2–3, pp. 77– 86, May 2008.

[34] R. C. Hunter and T. J. Beveridge, “High-resolution visualization of Pseudomonas aeruginosa PAO1 biofilms by freeze-substitution transmission electron microscopy,” J. Bacteriol., vol. 187, no. 22, pp. 7619–7630, Nov. 2005.

[35] B. K. Pierson and R. W. Castenholz, “Bacteriochlorophylls in gliding filamentous prokaryotes from hot springs,” Nat. New Biol., vol. 233, no. 35, pp. 25–27, Sep. 1971.

[36] S. Hanada, A. Hiraishi, K. Shimada, and K. Matsuura, “Chloroflexus aggregans sp. nov., a filamentous phototrophic bacterium which forms dense cell aggregates by active gliding movement,” Int. J. Syst. Bacteriol., vol. 45, no. 4, pp. 676–681, Oct. 1995.

[37] S. Hanada, K. Shimada, and K. Matsuura, “Active and energy-dependent rapid formation of cell aggregates in the thermophilic photosynthetic bacterium Chloroflexus aggregans,” FEMS Microbiol. Lett., vol. 208, no. 2, pp. 275–279, Mar. 2002.

[38] L. Cai et al., “Tad pilus-mediated twitching motility is essential for DNA uptake and survival of Liberibacters,” PLoS One, vol. 16, no. 10, p. e0258583, Oct. 2021.

[39] S. Yoshihara et al., “Mutational analysis of genes involved in pilus structure, motility and transformation competency in the unicellular motile cyanobacterium Synechocystis sp. PCC 6803,” Plant Cell Physiol., vol. 42, no. 1, pp. 63–73, Jan. 2001.

[40] Y. Yang et al., “Phototaxis in a wild isolate of the cyanobacterium Synechococcus elongatus,” Proc. Natl. Acad. Sci. U. S. A., vol. 115, no. 52, pp. E12378–E12387, Dec. 2018.

[41] R. Allen, B. E. Rittmann, and R. Curtiss 3rd, “Axenic Biofilm Formation and Aggregation by Synechocystis sp. Strain PCC 6803 Are Induced by Changes in Nutrient Concentration and Require Cell Surface Structures,” Appl. Environ. Microbiol., vol. 85, no. 7, Apr. 2019, doi: 10.1128/AEM.02192-18.

[42] M. D. M. Aguilo-Ferretjans et al., “Pili allow dominant marine cyanobacteria to avoid sinking and evade predation,” Nat. Commun., vol. 12, no. 1, p. 1857, Mar. 2021.

[43] D. Schatz et al., “Self-suppression of biofilm formation in the cyanobacterium Synechococcus elongatus,” Environ. Microbiol., vol. 15, no. 6, pp. 1786–1794, Jun. 2013.

[44] N. C. Caiazza, R. M. Q. Shanks, and G. A. O’Toole, “Rhamnolipids modulate swarming motility patterns of Pseudomonas aeruginosa,” J. Bacteriol., vol. 187, no. 21, pp. 7351– 7361, Nov. 2005.

[45] T. Ursell, R. M. W. Chau, S. Wisen, D. Bhaya, and K. C. Huang, “Motility enhancement through surface modification is sufficient for cyanobacterial community organization during phototaxis,” PLoS Comput. Biol., vol. 9, no. 9, p. e1003205, Sep. 2013.

[46] N. Luo, S. Wang, J. Lu, X. Ouyang, and L. You, “Collective colony growth is optimized by branching pattern formation in Pseudomonas aeruginosa,” Mol. Syst. Biol., vol. 17, no. 4, p. e10089, Apr. 2021.

[47] B. R. Wucher, J. B. Winans, M. Elsayed, D. E. Kadouri, and C. D. Nadell, “Breakdown of clonal cooperative architecture in multispecies biofilms and the spatial ecology of predation,” Proc. Natl. Acad. Sci. U. S. A., vol. 120, no. 6, p. e2212650120, Feb. 2023.

[48] J. B. Winans, B. R. Wucher, and C. D. Nadell, “Multispecies biofilm architecture determines bacterial exposure to phages,” PLoS Biol., vol. 20, no. 12, p. e3001913, Dec. 2022.

[49] D. Nishiguchi, K. H. Nagai, H. Chaté, and M. Sano, “Long-range nematic order and anomalous fluctuations in suspensions of swimming filamentous bacteria,” Phys Rev E, vol. 95, no. 2–1, p. 020601, Feb. 2017.

[50] C. Tamulonis and J. Kaandorp, “A model of filamentous cyanobacteria leading to reticulate pattern formation,” Life, vol. 4, no. 3, pp. 433–456, Sep. 2014.

[51] H. Jeckel, et al., “Multispecies phase diagram of biofilm architectures reveals biophysical principles of biofilm development,” bioRxiv, p. 2021.08.06.455416, Aug. 15, 2022. doi: 10.1101/2021.08.06.455416.

[52] T. Xu et al., “Characterization of Mixed-Species Biofilms Formed by Four Gut Microbiota,” Microorganisms, vol. 10, no. 12, Nov. 2022, doi: 10.3390/microorganisms10122332.

[53] D. Ren, J. S. Madsen, S. J. Sørensen, and M. Burmølle, “High prevalence of biofilm synergy among bacterial soil isolates in cocultures indicates bacterial interspecific cooperation,” ISME J., vol. 9, no. 1, pp. 81–89, Jan. 2015.

[54] L. L. Wong et al., “Surface-layer protein is a public-good matrix exopolymer for microbial community organisation in environmental anammox biofilms,” ISME J., vol. 17, no. 6, pp. 803–812, Jun. 2023.

[55] Y.-M. Kim et al., “Diel metabolomics analysis of a hot spring chlorophototrophic microbial mat leads to new hypotheses of community member metabolisms,” Front. Microbiol., vol. 6, p. 209, Apr. 2015.

[56] J. Z. Lee et al., “Fermentation couples Chloroflexi and sulfate-reducing bacteria to Cyanobacteria in hypersaline microbial mats,” Front. Microbiol., vol. 5, p. 61, Feb. 2014.

[57] S.-I. Fukushima, “Analysis of gliding motility of the filamentous bacterium Chloroflexus aggregans.” [Online]. Available: https://core.ac.uk/download/pdf/235009017.pdf

[58] S. M. Boomer, K. L. Noll, G. G. Geesey, and B. E. Dutton, “Formation of multilayered photosynthetic biofilms in an alkaline thermal spring in Yellowstone National Park, Wyoming,” Appl. Environ. Microbiol., vol. 75, no. 8, pp. 2464–2475, Apr. 2009.

[59] B. M. Bebout and F. Garcia-Pichel, “UV B-Induced Vertical Migrations of Cyanobacteria in a Microbial Mat,” Appl. Environ. Microbiol., vol. 61, no. 12, pp. 4215–4222, Dec. 1995.

[60] X. Zhang, B. Ji, J. Tian, and Y. Liu, “Development, performance and microbial community analysis of a continuous-flow microalgal-bacterial biofilm photoreactor for municipal wastewater treatment,” J. Environ. Manage., vol. 338, p. 117770, Jul. 2023.

[61] M. Nierychlo, A. Milobedzka, F. Petriglieri, B. McIlroy, P. H. Nielsen, and S. J. McIlroy, “The morphology and metabolic potential of the Chloroflexi in full-scale activated sludge wastewater treatment plants,” FEMS Microbiol. Ecol., vol. 95, no. 2, Feb. 2019, doi: 10.1093/femsec/fiy228.

[62] A. Sood, N. Renuka, R. Prasanna, and A. S. Ahluwalia, “Cyanobacteria as Potential Options for Wastewater Treatment,” in Phytoremediation: Management of Environmental Contaminants, Volume 2, A. A. Ansari, S. S. Gill, R. Gill, G. R. Lanza, and L. Newman, Eds., Cham: Springer International Publishing, 2015, pp. 83–93.

[63] D. Brockmann, Y. Gérand, C. Park, K. Milferstedt, A. Hélias, and J. Hamelin, “Wastewater treatment using oxygenic photogranule-based process has lower environmental impact than conventional activated sludge process,” Bioresour. Technol., vol. 319, p. 124204, Jan. 2021.

[64] M. Nierychlo et al., “Candidatus Amarolinea and Candidatus Microthrix Are Mainly Responsible for Filamentous Bulking in Danish Municipal Wastewater Treatment Plants,” Front. Microbiol., vol. 11, p. 1214, Jun. 2020.

[65] K. Heimann, “Novel approaches to microalgal and cyanobacterial cultivation for bioenergy and biofuel production,” Curr. Opin. Biotechnol., vol. 38, pp. 183–189, Apr. 2016.

[66] M. Bozan, A. Schmid, and K. Bühler, “Evaluation of self-sustaining cyanobacterial biofilms for technical applications,” Biofilms, vol. 4, p. 100073, Dec. 2022.

[67] D. Bhaya et al., “Population level functional diversity in a microbial community revealed by comparative genomic and metagenomic analyses,” ISME J., vol. 1, no. 8, pp. 703–713, Dec. 2007.

[68] R. W. Castenholz, “Isolation and Cultivation of Thermophilic Cyanobacteria,” in The Prokaryotes: A Handbook on Habitats, Isolation, and Identification of Bacteria, M. P. Starr, H. Stolp, H. G. Trüper, A. Balows, and H. G. Schlegel, Eds., Berlin, Heidelberg: Springer Berlin Heidelberg, 1981, pp. 236–246.

[69] S. Hanada, A. Hiraishi, K. Shimada, and K. Matsuura, “Isolation of *Chloroflexus aurantiacus* and related thermophilic phototrophic bacteria from Japanese hot springs using an improved isolation procedure,” J. Gen. Appl. Microbiol., vol. 41, no. 2, pp. 119–130, 1995.

[70] J. Schindelin et al., “Fiji: an open-source platform for biological-image analysis,” Nat. Methods, vol. 9, no. 7, pp. 676–682, Jun. 2012.

[71] D. Ershov et al., “TrackMate 7: integrating state-of-the-art segmentation algorithms into tracking pipelines,” Nat. Methods, vol. 19, no. 7, pp. 829–832, Jul. 2022.

[72] R Core Team, “R: A Language and Environment for Statistical Computing,” 2024, R Foundation for Statistical Computing, Vienna, Austria. [Online]. Available: https://www.R-project.org/

[73] P. A. Zaini, L. De La Fuente, H. C. Hoch, and T. J. Burr, “Grapevine xylem sap enhances biofilm development by Xylella fastidiosa,” FEMS Microbiol. Lett., vol. 295, no. 1, pp. 129– 134, Jun. 2009.

[74] M. Román-Écija et al., “Two Xylella fastidiosa subsp. multiplex Strains Isolated from Almond in Spain Differ in Plasmid Content and Virulence Traits,” Phytopathology, vol. 113, no. 6, pp. 960–974, Jun. 2023.

[75] J. R. Kremer, D. N. Mastronarde, and J. R. McIntosh, “Computer visualization of three- dimensional image data using IMOD,” J. Struct. Biol., vol. 116, no. 1, pp. 71–76, Jan-Feb 1996.

## References

1. K. Onai, M. Morishita, T. Kaneko, S. Tabata, M. Ishiura, Natural transformation of the thermophilic cyanobacterium Thermosynechococcus elongatus BP-1: a simple and efficient method for gene transfer. Mol. Genet. Genomics 271, 50–59 (2004).

2. L. P. Sáez, et al., Cyanate Assimilation by the Alkaliphilic Cyanide-Degrading Bacterium Pseudomonas pseudoalcaligenes CECT5344: Mutational Analysis of the cyn Gene Cluster. Int. J. Mol. Sci. 20 (2019).

3. F. Bunbury, et al., Differential Phototactic Behavior of Closely Related Cyanobacterial Isolates from Yellowstone Hot Spring Biofilms. Appl. Environ. Microbiol. 88, e0019622 (2022).

